# POMC neurons functional heterogeneity relies on mTORC1 signaling

**DOI:** 10.1101/2020.03.25.007765

**Authors:** Nicolas Saucisse, Wilfrid Mazier, Vincent Simon, Elke Binder, Caterina Catania, Luigi Bellocchio, Roman A. Romanov, Isabelle Matias, Philippe Zizzari, Stephane Leon, Carmelo Quarta, Astrid Cannich, Kana Meece, Delphine Gonzales, Samantha Clark, Julia M. Becker, Giles S.H. Yeo, Florian T. Merkle, Sharon L. Wardlaw, Tibor Harkany, Federico Massa, Giovanni Marsicano, Daniela Cota

## Abstract

Hypothalamic Pro-opiomelanocortin (POMC) neurons are classically known to trigger satiety. However, they encompass heterogeneous subpopulations whose functions are unknown. Here we show that POMC neurons releasing GABA, glutamate or both neurotransmitters possess distinct spatial distribution, molecular signatures and functions. Functional specificity of these subpopulations relies on the energy sensor mechanistic Target of Rapamycin Complex 1 (mTORC1), since pharmacological blockade of mTORC1, by mimicking a cellular negative energy state, simultaneously inhibited POMC/glutamatergic and activated POMC/GABAergic neurons. Chemogenetics and conditional deletion of mTORC1 then demonstrated that mTORC1 blockade in POMC neurons causes hyperphagia. This is due to decreased POMC-derived anorexigenic *α*-melanocyte-stimulating hormone and the recruitment of POMC/GABAergic neurotransmission, which is restrained by cannabinoid type 1 receptor signaling. Genetic inhibition of glutamate release from POMC neurons also produced hyperphagia, recapitulating the phenotype caused by mTORC1 blockade. Altogether, these findings pinpoint the molecular mechanisms engaged by POMC neurons to oppositely control feeding, thereby challenging conventional views about their functions.

## **1.** Introduction

Food intake is a highly complex behavior controlled by the coordinated action of specialized cell types organized in specific networks. Within the arcuate nucleus (ARC) of the hypothalamus, two neuronal populations respectively expressing Pro-opiomelanocortin (POMC) and Agouti-related peptide (AgRP) oppositely control food intake due to antagonistic effects on the central melanocortin receptors type 4 (MC4R) (Krashes et al., 2016).

Both POMC and AgRP neurons respond to nutrient and hormonal signals reflecting short- and long-term changes in energy availability (Cota et al., 2007; Krashes et al., 2016). Energy deficit activates AgRP neurons, which drive food seeking and intake through the release of the MC4R antagonist AgRP, the Neuropeptide Y (NPY), and the inhibitory neurotransmitter *γ*-aminobutyric acid (GABA) (Fan et al., 1997; Tong et al., 2008). Conversely, energy surfeit activates POMC neurons, which typically promote satiety by releasing the MC4R agonist *α*-melanocyte-stimulating hormone (*α*-MSH) (Krashes et al., 2016). Short-term food consumption after a prolonged fast (also known as refeeding) increases the activity of POMC neurons, thereby regulating meal size through *α*-MSH dependent activation of MC4R in the hypothalamic paraventricular nucleus (PVN), among other brain areas (Balthasar et al., 2005; Singru et al., 2007; Fekete et al., 2012). Besides, recent studies have shown that sensory detection of food occurring before actual food intake simultaneously inhibits AgRP neurons and activates POMC neurons, thereby preparing the organism to the ingestion of food (Brandt et al., 2018; Chen et al., 2015). Thus, the classic view of AgRP and POMC neurons as “Yin and Yang” partners regulating food intake through their opposite action on MC4R may be an over-simplistic and partial depiction of the relevant circuit. Indeed, AgRP neurons seem physiologically relevant for the search of food rather than food intake, while POMC neurons may actually be the ones in charge of controlling the whole feeding sequence, from its start to its termination, as suggested by recently published evidence (Brandt et al., 2018; Chen et al., 2015). In support of this hypothesis, POMC cleavage does not only produce *α*-MSH, but also the opioid β-endorphin (*β*-EP), which stimulates food intake (Koch et al., 2015; Wardlaw, 2011). Besides, and differently from AgRP neurons, which are almost exclusively GABAergic (Tong et al., 2008), POMC neurons include GABAergic and glutamatergic cells, as well as neurons containing both these neurotransmitters (Dicken et al., 2012; Hentges et al., 2004; Hentges et al., 2009; Mazier et al., 2019; Wittmann et al., 2013). Finally, recent single cell transcriptomic studies have revealed that POMC neurons are molecularly heterogeneous (Campbell et al., 2017; Henry et al., 2015; Lam et al., 2017). Yet the specific roles of molecularly identified POMC neurons subpopulations in the context of the regulation of food intake remain unexplored.

The mechanistic target of rapamycin (mTOR) kinase is a critical cellular energy sensor, whose increased activity is a readout of increased energy availability (Haissaguerre et al., 2014; Saxton and Sabatini, 2017). By forming two distinct complexes (mTOR complex 1 - mTORC1, and mTOR complex 2 - mTORC2), mTOR controls cellular metabolism in response to nutrients, growth factors, hormones and cellular stress (Haissaguerre et al., 2014; Saxton and Sabatini, 2017). In neurons, the mTOR pathway regulates soma size, dendrite and axon growth (Bockaert and Marin, 2015), and it affects both glutamatergic and GABAergic transmission (Weston et al., 2012). Others and we have shown that mTORC1 signaling participates in the regulation of energy balance by modulating the function of AgRP and POMC neurons (Burke et al., 2017; Cota et al., 2006; Dagon et al., 2012; Smith et al., 2015;). mTORC1 also regulates oxidative metabolism in POMC neurons (Haissaguerre et al., 2018) and glutamatergic transmission in the PVN (Mazier et al., 2019). However, it is unknown whether mTORC1 signaling has a role in matching energy availability to POMC neuronal function depending on POMC neurotransmitter subtypes, and how this may impact food intake.

GABAergic and glutamatergic transmission, including in brain areas targeted by POMC neurons, is regulated by specific mechanisms (Dietrich and Horvath, 2013). Among these, the retrograde suppression of neurotransmitter release through endocannabinoids acting at the presynaptic cannabinoid type 1 receptor (CB_1_R) is well described (Busquets-Garcia et al., 2018; Castillo et al., 2012). Brain CB_1_R activation generally stimulates food intake (Mazier et al., 2015) and it offsets *α*-MSH effects in the PVN (Mazier et al., 2019; Monge-Roffarello et al., 2014; Verty et al., 2004). However, the impact of brain CB_1_R on food intake is also cell-type specific. For instance, while CB_1_R-dependent inhibition of glutamatergic transmission induces hyperphagia, CB_1_R-depedent inhibition of GABAergic neurons is hypophagic (Bellocchio et al., 2010; Soria-Gomez et al., 2014a). Whether CB_1_R on POMC neurons might affect feeding by participating to the modulation of neurotransmitter release is not known. Here we have used neuroanatomical, transcriptomic, pharmacological, genetic, chemogenetic, electrophysiological, optogenetic and behavioral approaches to address the physiological role of POMC neurons in the control of food intake. These experiments have revealed functionally distinct POMC neuronal subclasses, whose activity is under the control of mTORC1 and CB_1_R signaling. These findings pinpoint the molecular mechanisms engaged by different POMC neurons subpopulations to oppositely regulate feeding, and suggest that one physiological purpose of the mTORC1 pathway is to coordinate both neuropeptidergic- and neurotransmitter-dependent responses of POMC neurons in relation to energy availability. Altogether, this evidence challenges classic notions about the role of POMC neurons as exclusive drivers of satiety.

## 2. Material and Methods

### 2.1. Study Approval

All experiments were conducted in strict compliance with the European Union recommendations (2013/63/EU) and were approved by the French Ministry of French Ministry of Higher Education, Research and Innovation (animal experimentation authorization n°3309004) and the local ethical committee of the University of Bordeaux (DIR1354; APAFIS13232). For the triple FISH studies (see further below), no regulated procedures under the Animals (Scientific Procedures) Act 1986 were carried out on the animals used and the studies was reviewed by the University of Cambridge Animal Welfare and Ethical Review Body (AWERB).

### 2.2. Animals

Male mice, aged 7-16 weeks, were housed individually in standard plastic rodent cages, maintained on a 12-h light-dark cycle (lights off at 1300 h) with *ad libitum* access to pelleted chow (A03, SAFE, France) and water, unless otherwise specified. Number of animals for each experiment is detailed in the figure legends.

C57BL/6J mice (Janvier, France or Charles River, UK), POMC-Cre mice [Tg(Pomc1-cre)16Lowl/J, JAX Stock #005965, The Jackson Laboratory, USA], POMC-YFP mice, POMC-*Rptor*-KO, POMC-*Rictor*-KO, POMC-*CB_1_*-KO, POMC-CreER^T2^/vGlut2-flox (thereafter called POMC-*vGlut2*-KO after tamoxifen administration), POMC-CreER^T2^-Ai6 and their control littermates were used.

POMC-YFP mice were obtained by crossing POMC-Cre mice with Rosy mice [B6.129X1-Gt(ROSA)26Sortm1(EYFP)Cos/J, JAX Stock #006148, The Jackson Laboratory]. POMC-*Rptor*-KO were generated by crossing POMC-Cre mice with Rptor-Flox mice (B6.Cg-Rptortm1.1Dmsa/J, JAX Stock #013188). POMC-*Rictor*-KO were obtained by crossing POMC-Cre mice with Rictor-Flox mice (Rictortm1.1Klg/SjmJ, JAX Stock #020649). POMC-*CB_1_*-KO were generated by crossing POMC-Cre mice with CB_1_-Flox mice (Marsicano et al., 2003). Tamoxifen inducible POMC-*vGlut2*-KO were generated by crossing POMC-CreER^T2^ mice [C57BL/6J-Tg(Pomc-creERT2), kind gift of Dr. Joel Elmquist at UT Southwestern, (Berglund et al., 2013)] with vGlut2-flox mice (Slc17a6tm1Lowl/J, JAX stock #012898). Inducible POMC-CreER^T2^-Ai6 mice were generated by crossing POMC-CreER^T2^ mice with Ai6 mice (B6.Cg-Gt(ROSA)26Sortm6(CAG-ZsGreen1)Hze/J, JAX stock #007906). New conditional KO lines were generated following a three step backcrossing method (Bellocchio et al., 2013). All lines were in a mixed genetic background. All animals used in experiments involving mutant mice were littermates. We have previously characterized POMC-*Rptor*-KO mice and POMC-*CB_1_*-KO mice (Haissaguerre et al., 2018; Mazier et al., 2019). Effective Cre-mediated deletion of rictor in POMC neurons was assessed by immunohistochemistry. Appropriate recombination after tamoxifen administration was evaluated by performing immunohistochemistry in POMC-CreER^T2^-Ai6 mice, and by assessing vGlut2 deletion in POMC neurons by PCR (see below).

### 2.3. PCR for mouse genotyping

PCR on tail biopsies or hypothalamic samples (to assess vGlut2 excision in POMC-*vGlut2*-KO mice) was performed as in (Bellocchio et al., 2013) by using specific primers (see Supplementary Table S1 in Supplementary Information).

### 2.4. AAV vectors for hM3Dq and channelrhodopsin-2 (ChR2) expression

Gq-coupled human M3 muscarinic DREADD (hM3Dq) fused to mCherry (Alexander et al., 2009), provided by Brian L. Roth (University of North Carolina, Chapel Hill, NC, USA) was subcloned in a CAG-DIO rAAV vector for Cre-dependent expression (Atasoy et al., 2008) using standard molecular biology techniques. AAV-DIO-hChR2-mCherry was generated using pAAV-Ef1a-DO-hChR2(H134R)-mCherry-WPRE-pA plasmid as backbone (ADDGENE #37082). Vectors used were of an AAV1/AAV2 mixed serotype, and were generated by calcium phosphate transfection of HEK-293T cells and subsequent purification as described (Monory et al., 2006).

### 2.5. Tamoxifen treatment for the induction of Cre-mediated recombination

Corn oil (Sigma-Aldrich, France), pre-heated to 55°C, was added to 1g/mL suspension of tamoxifen (Sigma-Aldrich) in 100% ethanol (VWR, France) obtain a 50mg/mL stock solution. Mice received daily 150mg/kg of tamoxifen for 5 consecutive days by oral gavage (3mL/kg) done using polypropylene flexible gavage needles (Instechlabs, USA).

### 2.6. Stereotaxic surgery

Mice were anesthetized with a mixture of ketamine (100 mg/kg, ip) and xylazine (10 mg/kg, ip), placed in a stereotaxic holder (David Kopf Instruments, USA) and implanted with a cannula in the lateral cerebral ventricle (coordinate AP/DV/ML = −0.5/−2.1/-1.2 mm). Cannula placement was verified by administering 5 µg of neuropeptide Y (NPY) (Phoenix Pharmaceuticals Inc., France) in 0.1 M PBS (pH 7.4) and assessing subsequent food intake (Brown et al., 2006).

#### 2.6.1. Stereotaxic virus injection

AAV vectors (500 nL) were bilaterally infused into the ARC (coordinate AP/DV/ML = −1.5/−5.5/±0.2 mm; infusion speed = 100 nL/min; UltraMicroPump UMP3 (one) with SYS-Micro4 Controller, WPI, USA) using a Hamilton syringe with NanoFil needle (85 μm tip diameter). AAV-DIO-hM3Dq-mCherry vectors were injected into the ARC of POMC-Cre^+/+^ (thereafter named POMC-ARC^hM3Dq^ mice) and of POMC-Cre^-/-^ mice (thereafter named POMC-ARC^hM3Dq^-control mice). Mice were then implanted with cannulae into the lateral ventricle as detailed above. Mice were used for the behavioral studies after at least 4 weeks from the intra-ARC AAV administration. For optogenetic electrophysiology studies, 5-weeks old POMC-Cre^+/+^ mice were stereotaxically and bilaterally injected with an AAV-DIO-hChR2-mCherry virus and used for electrophysiology 4-5 weeks after the intra-ARC AAV administration.

### 2.7. Drugs preparations and food intake studies

Rapamycin (RAPA, EMD Millipore, USA), and picrotoxin (Ptx, Sigma-Aldrich), were dissolved in DMSO, the *α*-MSH analog melanotan II (MTII, Phoenix Pharmaceuticals, USA) was dissolved in phosphate-buffered saline (PBS), and clozapine-N-oxide (CNO, Sigma-Aldrich) was dissolved in saline.

#### 2.7.1. Recording of basal food intake

Before undergoing pharmacological studies, 24h basal, unstimulated, food intake was recorded over several days in POMC-*Rictor*-KO, POMC-*CB_1_*-KO, POMC-*vGlut2*-KO mice and their control littermates.

#### 2.7.2. Fasting-induced food intake studies

24h fasted mice received an icv injection of RAPA (25 µg in 1 µl DMSO) or its vehicle, which were administered 4h before dark onset. The same protocol was followed when Ptx (0.03 µg in 1 µl DMSO), MTII (0.02 µg in 1 µl PBS) or CNO (1 mg/kg in saline, ip) were combined with RAPA. Ptx or MTII were simultaneously administered icv with RAPA; CNO or its vehicle were administered 15 min before the icv administration of RAPA or its vehicle. Food was returned immediately after the drugs injections. Whenever possible the same animals belonging to the same genotype underwent the different treatments. Animals showing sign of stress or malaise during the experiments were removed from further analysis.

#### 2.7.3. MTII and Ptx dose-responses

The MTII and the Ptx dose-response were carried out in 24h fasted C57BL/6J mice that received MTII (0.5, 0.25, 0.125, 0.0625, 0.03, 0.02 or 0.01 µg/µL in saline, icv) or Ptx (0.03, 0.06 or 0.09 µg/µL in DMSO, icv) respectively 1 h and 4 h before dark onset. Food was returned immediately after the drugs injections.

#### 2.7.4. Palatable food intake study

C57Bl/6J male mice were singly housed and divided into two body weight-matched groups called “chow” and “palatable”. While the chow group continued to eat chow (A03, SAFE, France), the palatable group was switched from chow to a palatable diet (D12492, Research Diets Inc.) during the light phase. Two hours after the switch, food intake in both groups was measured and mice underwent anesthesia and perfusion for subsequent collection of brains for neuroanatomical analysis.

### 2.8. Immunohistochemistry (IHC)

Mice were deeply anesthetized using pentobarbital given ip and then perfused transcardially with ice-cold PBS, pH 7.4, followed by 4% paraformaldehyde (PFA, Sigma-Aldrich) in PBS with 0.2% picric acid. Brains were extracted and postfixed in 4% PFA overnight at 4°C, then cryoprotected with 30% sucrose in PBS at 4°C. Coronal sections (30 μm) were cut with a cryostat (CM1950, Leica, Germany), collected in PBS and stored in antifreeze solution (30% ethylene glycol, 30% glycerol in KPBS) at −20°C until further used.

#### 2.8.1. Single-labeling IHC

Brain sections from 90 min refed POMC-ARC^hM3Dq^ mice were processed for the co-localization of mCherry and POMC. Sections were first incubated with 10% normal goat serum (Dako, Denmark) and then with rabbit anti-POMC antibody (1:2000; Phoenix Pharmaceuticals) overnight at 4°C. The next day, sections were washed in PBS and incubated for 1h with A488-conjugated secondary goat anti-rabbit antibody (1:500, Cell Signaling, USA). Similar procedures were followed for POMC labelling in brain sections from POMC-CreER^T2^-Ai6 mice. Because of the endogenous green fluorescence of Zsgreen1expressed in this mouse line after tamoxifen administration, an A647-conjugated secondary donkey anti-rabbit antibody (1:500, Cell Signaling) was used to reveal the POMC staining.

#### 2.8.2. Double-labeling IHC

Brain sections from POMC-*Rictor*-KO and their control littermates were processed for the co-localization of rictor and POMC. Sections were first incubated with 10% normal goat serum (Dako) and then with rabbit anti-rictor antibody (1:1000, Bethyl Laboratories, Inc., USA) overnight at 4°C. The next day sections were washed in PBS and incubated for 1h with A647-conjugated secondary goat anti-rabbit antibody (1:500, Cell Signaling). Sections were washed in PBS and incubated with Goat Fab Fragment Anti-Rabbit IgG (1:100, Jackson ImmunoResearch Laboratories, USA) to avoid cross-reactivity. Sections were then washed in PBS and blocked with 10% normal goat serum (Dako) and incubated with rabbit anti-POMC antibody (1:2000; Phoenix Pharmaceuticals) overnight at 4°C. The next day sections were washed in PBS and incubated for 1h with A488-conjugated secondary goat anti-rabbit antibody (1:500, Cell Signaling).

#### 2.8.3. Evaluation of chemogenetic activation of POMC neurons

Free-fed POMC-ARC^hM3Dq^ mice received a single injection of CNO (1 mg/kg, ip) or its vehicle 90 minutes before sacrifice and expression of c-Fos and phosphorylated S6 ribosomal protein (p-S6) in mCherry-expressing POMC neurons was evaluated. Collected sections were first incubated with 10% normal donkey serum (Jackson ImmunoResearch Laboratories) and then with goat anti-c-Fos antibody (1:250; Santa Cruz Biotechnology, USA) overnight at 4°C. The next day sections were washed in PBS and incubated for 1h with A488-conjugated secondary donkey anti-goat antibody (1:500, EMD Millipore, USA). Sections were then washed in PBS and blocked with 10% normal goat serum (Dako) and subsequently incubated with rabbit anti-p-S6 ser240/244 antibody (1:200; Cell Signaling) overnight at 4°C. The next day sections were washed in PBS and incubated for 1h with A647-conjugated secondary goat anti-rabbit antibody (1:500, Cell Signaling).

#### 2.8.4. Triple-labeling IHC

Brain sections from 24-h fasted and 90 min refed C57BL/6J mice icv treated with RAPA or its vehicle were processed for the expression of c-Fos and p-S6 in POMC neurons. Sections were first incubated with 10% normal donkey serum (Jackson ImmunoResearch Laboratories) and then with goat anti-c-Fos antibody (1:250; Santa Cruz Biotechnology) overnight at 4°C. The next day sections were washed in PBS and incubated for 1 h with A594-conjugated secondary donkey anti-goat antibody (1:500, EMD Millipore). Sections were then washed in PBS and blocked with 10% normal goat serum (Dako) and then incubated with rabbit anti-p-S6 antibody (1:200; Cell Signaling) overnight at 4°C. The next day sections were washed in PBS and incubated for 1 h with biotinylated goat anti-rabbit IgG secondary antibody (1:400, Vector Labs), followed by washes and 1 h of amplification with an avidin-biotin complex (Vector Labs). The sections were then washed and incubated with a FITC-conjugated streptavidin (1:400, Vector Labs). Finally, sections were washed in PBS and incubated with Goat Fab Fragment Anti-Rabbit IgG (1:100, Jackson ImmunoResearch Laboratories) to avoid cross-reactivity. Sections were then washed in PBS and blocked with 10% normal goat serum (Dako) and then incubated with rabbit anti-POMC antibody (1:2000, Phoenix Pharmaceuticals) overnight at 4°C. The next day sections were washed in PBS and incubated for 1 h with A647-conjugated secondary goat anti-rabbit antibody (1:500, Cell Signaling). Sections were then mounted and cover-slipped.

#### 2.8.5. Quantification of the immunohistochemical signal

Fluorescent images were acquired with a confocal microscope (SP8-STED, Leica, Germany), corrected for brightness and contrast, and analyzed using ImageJ (National Institutes of Health, USA).

### 2.9. Fluorescent in situ hybridization (FISH)

#### 2.9.1. Double FISH

Brain sections from perfused POMC-CreER^T2^-Ai6 mice were used to assess possible co-expression of AgRP and POMC mRNA. Digoxigenin-labeled riboprobe against mouse AgRP (AgRP-DIG, Allen Brain Atlas, #RP_050419_04_D06) and fluorescein-labeled probe against mouse POMC (POMC-FITC, Allen Brain Atlas, #RP_Baylor_102974) were prepared as in (Marsicano et al., 2002). Free-floating sections were treated with 0.2M HCl then acetylated with 0.25% acetic anhydride in 0.1 M triethanolamine, pH=8.0 for 10 min. Between all steps, sections were rinsed in PBS with 0.01% diethylpyrocarbonate. AgRP-DIG and POMC-FITC probes were dissolved 1/1000 in hybmix solution and heated at 90°C for 5min. After hybridization overnight at 70°C, sections were washed at 65°C with increased stringency buffers (5X SSC for 5min, 2X SSC+50%Formamide Amide, 1X SSC+50%Formamide Amide and 0.1X SSC for 30min each, with 0.1% Tween20 added to each buffer). After blocking 1h in TNB buffer (AkoyaBiosciences, prepared as per manufacturer’s instructions), sections were incubated overnight at 4°C in HRP-anti-DIG antibody (1/1500, Sigma) in TNB. The revelation was made using TSA-Cy3 kit (1/100, 30min, AkoyaBiosciences). After quenching the peroxidase with 3% H2O2 in PBS (30min) and 0.2M HCl treatment (20min), sections were incubated in HRP-anti-FITC antibody (1/1500, Sigma) in TNB for 2h and the revelation made using TSA-FITC kit (1/100, 30min, AkoyaBiosciences).

#### 2.9.2. Triple FISH

C57BL/6J mice were killed by an overdose of pentobarbital. The brains were removed, instantly embedded in OCT Embedding Medium (Thermo Scientific) and stored at −80°C. 10 µm thick coronal sections of the entire ARC were prepared on a Leica CM1950 Cryostat. One section every 100 µm was stained and analyzed from each animal. The following primers were used for FISH antisense probe production: Pomc forward GCAGTGACTAAGAGAGGCCACT, Pomc reverse ATTTAGGTGACACTATAGAAGAGGACTGCCATCTCCCCACAC; Gad67 forward GAAAGGGCCAATTCAGTCAC, Gad67 reverse ATTTAGGTGACACTATAGAAGAGCTGCCTTCAGTGAGATGGCCTAG; vGlut2 forward TCATTGCTGCACTCGTCCACTA, vGlut2 reverse ATTTAGGTGACACTATAGAAGAGCCCTGGGATAGTTTGCAGTCCA. Reverse primers included a SP6 recognition sequence. A hypothalamic mouse RNA library was reverse transcribed into a cDNA library and DNA templates for the relevant transcripts were obtained via PCR using the above primers. Subsequently, these templates were in vitro transcribed into antisense RNA probes using MAXIscript SP6 In Vitro Transcription Kit (Thermo Scientific). 35% of the UTP in each reaction was replaced with either Fluorescein-12-UTP (Roche), Digoxigenin-11-UTP (Roche) or DNP-11-UTP (PerkinElmer) for Pomc, VGlut2 and Gad67 probe production, respectively. For FISH, the following antibodies and tyramide solutions were used: Anti-Fluorescein-POD Fab fragments (Roche), Anti-Digoxigenin-POD Fab fragments (Roche) and Anti-DNP HRP Conjugate (PerkinElmer) as well as Cyanine 3 Tyramide Reagent and Cyanine 5 Tyramide Reagent and Fluorescein Tyramide Reagent (all PerkinElmer), respectively. Briefly, cryosections were fixed in 4% PFA, washed in PBST and 0.1 M TEA and acetylated, washed again in PBST, and in 5x SSC/50% formamide at 65°C. They were pre-hybridised in prehybridization buffer for 30 min at 65°C and incubated with hybridisation buffer containing all three antisense RNA probes overnight at 65°C. Following washes in 5x SSC, 1x SSC/50% formamide, 2x SSC and 0.2x SSC at 65°C, slides were rinsed with Maleic Acid Buffer with Tween20 (MABT), incubated with one of the above antibodies for 1h, washed with MABT again and developed with one of the above tyramide solutions for 20 min in the dark. Slides were washed with PBST and the remaining enzyme activity of the antibody quenched with 3% H2O2 in PBS for 30 min and subsequent citrate buffer at 95°C for 10 min. The antibody and tyramide incubation steps and the quenching procedure were repeated twice with the remaining antibodies and tyramide solutions. Finally, slides were washed several times in MABT, counterstained with DAPI in PBST and mounted with Aqua-Poly/Mount (Polysciences). All steps were carried out at room temperature unless stated otherwise.

#### 2.9.3. Image acquisition and analysis

Whole-slide images were acquired on an Axio Scan.Z1 (Zeiss). Image analysis and cell counts were obtained manually in the HALO software (Indica Labs Inc.). Analysis thresholds were applied before images were taken. Brightness and contrast were subsequently modified to facilitate signal visibility. A schematic image of each analyzed coronal brain section was created using Adobe Illustrator CS5 (Adobe Systems Software). For each POMC neuron, its distance to the ventricle and the subpopulations it belonged to was extracted from these schematic images by a custom-written Python script (Graham N. Stratton).

### 2.10. Triple-FISH combined with IHC

Free-floating brain sections from perfused brains of C57BL/6J mice having undergone the palatable diet test were used. Detailed description of the technical procedure used to combine FISH with IHC is included in the Supplementary Information. Quickly, after peroxidase quenching and biotin blocking, primary antibody anti-cFos was incubated overnight at 4°C. The next day, HRP-linked secondary antibody was incubated, followed by a Biotin-TSA amplification. After HRP quenching, TSA-deposited biotins were fixed by a 10min incubation in 4% formaldehyde. Sections were then pre-treated for the FISH (HCl and acetylation) and vGlut2-DIG, GAD65-FITC, GAD67-FITC and POMC-DNP probes were incubated in the hybmix overnight at 70°C. The probes used were as detailed earlier for vGlut2 and POMC (probe #RP_Baylor_102974), the GAD65 probe was a kind gift of Dr. Beat Lutz (Univ of Mainz, Germany), while for GAD67 we used RP_040324_01_F01 from the Allen Brain Atlas. The next day, after stringent washes, probes were sequentially revealed using anti-DIG, anti-FITC and anti-DNP secondary antibodies followed by Cy3-TSA, FITC-TSA and homemade Cy5-TSA amplification respectively. Between each probe revelation, HRP were quenched. Finally, biotins were revealed by ABC (Vector) followed by a Coumarin-TSA amplification. Sections were mounted using ProLong Gold medium and cover-slipped.

#### 2.10.1. Image acquisition and analysis

Stacks of images containing each hemi-ARC (left and right) were taken using a SP8 confocal microscope (Leica, Wetzlar, Germany). The voxel size was 0.284 x 0.284 x 0.7868 µm3, with a bit depth of 8 bits per pixels. Image analysis was performed using ImageJ. After brightness and contrast adjustment to reduce background, the co-expression of our different markers was assessed. For each image, we used the multi-point tool available in ImageJ to select and count POMC neurons that co-expressed (or not) c-Fos, vGlut2 and/or GAD65/67. Only one side (left of right) was quantified for each animal.

### 2.11. Hypothalamic neuropeptides assays

Hypothalami from 24h fasted or 2h refed C57BL/6J mice, treated icv with RAPA or its vehicle, were extracted in 0.1N HCl and assayed for *α*-MSH and *β*-EP by RIA as in (Savontaus et al., 2004). ACTH RIA was performed as in (Papadopoulos and Wardlaw, 1999). ELISA for POMC protein was performed with reagents provided by Dr. Anne White at the University of Manchester (Tsigos et al., 1993). Data are expressed versus mg of protein.

### 2.12. Endocannabinoid measurements

Hypothalamus, hippocampus and cortex of 24h fasted or 1h refed C57BL/6J mice, treated icv with RAPA or its vehicle, were dissected, flash frozen and stored at −80°C until analysis. In other experiments, hypothalami were collected from 1h refed C57BL/6J mice injected icv with MTII (0.02µg) or PBS, with RAPA or with RAPA combined to MTII. Finally, the mediobasal hypothalamus was collected from 1h refed POMC-ARC^hM3Dq^ mice having received CNO or its vehicle ip and RAPA icv. Measurements of AEA and 2-AG were performed as in (Gatta-Cherifi et al., 2012). Endocannabinoids content was expressed versus mg of tissue.

### 2.13. Single-cell RNA sequencing analysis

RNA-seq data on *Pomc*-expressing neurons (≥2 mRNA copies per cell) were extracted from our publicly available dataset on hypothalamic neurons obtained from 2-4 weeks old C57BL/6N mice (GEO accession number: GSE74672, (Romanov et al., 2017). We evaluated the distribution of cells expressing *Pomc* transcripts in the hypothalamus through public databases on in situ mRNA hybridization (Allen Brain Atlas, Mouse Brain, experiments 2493 and 2494) and noted highly specific *Pomc* expression in the ARC. This validates that *Pomc*-expressing neurons in our single-cell RNA-seq dataset originate from the ARC, and can be used for subtype deconvolution. For the validity of our following analysis, we checked cells for outliers using Principal Component Analysis to exclude those neurons coming to the dataset because of the contamination/ectopic *Pomc* gene transcription. Finally, we performed differential gene expression profiling of *Pomc*^+^ neurons by using Genepattern 3.9.8 (Reich et al., 2006); https://genepattern.broadinstitute.org) with the “ComparativeMarkerSelection” and “ExtractComparativeMarkerResults” modules. Data were then rendered into heatmaps (HeatMapImage module) for maximum visual clarity. The Database for Annotation, Visualization and Integrated Discovery (DAVID) Bioinformatics Resources 6.7 was used to identify the genes that had statistically significant differences in expression among the different POMC neuron subpopulations (Huang da et al., 2009). To interpret our data, we used the «Functional annotation chart» tool. This tool associates gene ID with a biological term which belongs to one out of the 40 annotation categories available in DAVID, including Gene Ontology terms and Kyoto Encyclopedia of Genes and Genomes (KEGG) pathways. This extended annotation coverage increases the analytic power by allowing investigators to analyze their genes from many different biological aspects in a single space. Each functional annotation was associated with an enrichment score, which depended on the distribution of the enrichment (p-values) of each gene. A good enrichment score was obtained when most of genes had good p-values. This score is a relative score instead of a statistical probability with a minimum and a maximum value. This means that enrichment scores could be considered only together, by comparing them.

### 2.14. Electrophysiology

#### 2.14.1. Brain slices preparation

Acute coronal brain slices were prepared from 8 to 12 week-old mice expressing YFP or Chr2 coupled to mCherry selectively in POMC neurons or from POMC-*CB_1_*-KO and their control littermates. Animals were anesthetized with isoflurane, the brain extracted and immediately placed in an ice-cold oxygenated cutting solution (in mM: 180 Sucrose, 26 NaHCO_3_, 11 Glucose, 2.5 KCl, 1.25 NaH_2_PO_4_, 12 MgSO_4_, 0.2 CaCl_2_, saturated with 95% O2-5% CO2). Slices were obtained using a vibratome (VT1200S Leica, Germany) and transferred into a 34°C bath of oxygenated aCSF (in mM: 123 NaCl, 26 NaHCO_3_, 11 Glucose, 2.5 KCl, 1.25 NaH_2_PO_4_, 1.3 MgSO_4_, 2.5 CaCl_2_; osmolarity 310 mOsm/l, pH 7.4) for 30 minutes and then cooled down progressively till room temperature (RT; 23-25°C) in oxygenated aCSF. After a 45 min recovery period at RT, slices were bisected by cutting along the third ventricle axis. The hemi-slice was anchored with platinum wire at the bottom of the recording chamber and continuously bathed in oxygenated aCSF (32-34°C; 2ml/min) during recording.

#### 2.14.2. Evaluation of WIN effect on mIPSC on PVN parvocellular neurons of POMC-*CB_1_* mice

Patch electrodes were pulled (micropipette puller P-97, Sutter instrument, USA) from borosilicate glass (O.D. 1.5 mm, I.D. 0.86 mm, Sutter Instrument) to a resistance of 2-4 m*Ω*. Electrophysiological data were recorded using a Multiclamp 700B amplifier (Molecular devices, UK), low-pass filtered at 4 kHz and digitized at 10Hz (current clamp) or 4 Hz (voltage clamp) (Digidata 1440A, Molecular devices, UK). Signals were analyzed offline (Clampfit software, pClamp 10, Molecular devices, UK). The pipette internal solution contained [in mM: 125 potassium gluconate, 5 KCl, 10 Hepes, 0.6 EGTA, 2 MgCl2, 7 Phosphocreatine, 3 adenosine-5’-triphosphate (magnesium salt), 0.3 guanosine-5’-triphosphate (sodium salt) (pH adjusted to 7.25 with KOH; osmolarity 300 mOsm/l adjusted with d-Mannitol; liquid junction potential −14.8mV corrected on the data presented)]. Parvocellular neurons were differentiated from magnocellular neurons immediately after patch rupture by preliminary electrophysiological analysis. Indeed, when submitted to a depolarizing current magnocellular neurons exhibit a large transient outward rectification, not found in parvocellular neurons (Luther and Tasker, 2000). Then cells were switched in voltage clamp (Vh = −70 mV). Miniature GABAergic transmission was pharmacologically isolated using a mix of NBQX (10 μM), APV (50 μM) and tetrodotoxin (TTX, 1μM). Note that as a control, few cells were perfused with a GABAA selective antagonist (picrotoxin, 100 μM) at the end of the experiment to fully block miniature transmission. To challenge the CB_1_ receptor, the selective agonist (WIN55-212, 5 µM) was added. For statistical analysis, miniature events frequency and amplitude during the last 4 minutes of baseline were compared to the same parameters after 10 minutes of drug perfusion (4 minutes also).

#### 2.14.3. Evaluation of RAPA effect on POMC neurons firing

Fluorescent POMC-YFP neurons were identified onto 250 μm thick slices obtained from 8-12 weeks old male POMC-YFP mice, using fluorescence/infrared light (pE-2 CoolLED excitation system, UK). Neurons action potential firing was monitored in whole-cell current-clamp recording configuration. For each recorded neuron, membrane potential (Em), membrane capacitance (Cm) and membrane resistance (Rm) were collected right after cell opening. After a 10 min baseline, RAPA (200 nM) was added in the bath for 20 minutes. For the analysis, considering the variability of the POMC neurons basal firing, the significance of RAPA-induced changes in action potential firing was calculated by comparing the firing rate before and during RAPA application (bin size of 10 sec) using normalization via Z-score transformation of individuals’ instantaneous frequencies values (Courtin et al., 2014). Z-score values were calculated by subtracting bin average instantaneous frequencies to the average baseline instantaneous frequency established over at least 120 sec preceding RAPA application and by dividing the difference by the baseline standard deviation. Scores under drug application were obtained comparing the firing rate of recorded neurons 120 sec before and after 4 min of drug application. Neurons with at least 5 significant positive or negative z-score bins (z-score > ±1.96, p<0.05) after RAPA application onset were considered as drug-responsive neurons.

#### 2.14.4. Effect of RAPA on light-evoked POMC to parvocellular neurons neurotransmission

350μm slices were prepared from brains of POMC-Cre^+/+^ mice bilaterally injected with an AAV-DIO-hChR2. The aCSF was modified in order to increase the neurotransmission success rate (3 mM Ca2+, 0.1 mM Mg2+ and 100μM 4 aminopyridine). Parvocellular neurons were switched in voltage clamp configuration (Vh = −70mV) and synaptic transmission was evoked with a 5 milliseconds flash of blue light (470nm, 1-3mW at 0.03 Hz; pE-2 CoolLED excitation system). Light-evoked glutamatergic or GABAergic transmission was pharmacologically isolated using picrotoxin (100μM) or a NBQX/APV mix (10μM/50μM), respectively. RAPA was added for 20 minutes after at least 10 minutes of stable amplitude light-evoked eEPSCs or eIPSCs. At the end of each experiment, responses were fully blocked using TTX (1μM). For statistical analysis, mean eEPSCs or eIPSCs amplitudes during the last 4 minutes of baseline period were compared to the amplitudes obtained in the same cells after 5 minutes of RAPA perfusion.

#### 2.14.5. POMC neuronal type characterization by IHC after recording

During recording POMC neurons were filled via passive diffusion of biocytin (0.4% in internal solution, Sigma, France). Afterwards, slices were fixed overnight at 4°C in 4% PFA-containing phosphate buffer (PB). After washing (3×15min in PB), slices were permeabilized and saturated for 3h at RT (0.3% triton, 10% bovine serum albumin in PB). Slices were incubated with primary antibody (anti-GAD65/67 made in rabbit, 1/2000, Millipore) for 72h at 4°C. Then the fluorophore-coupled secondary antibody (A647-conjugated secondary goat anti-rabbit antibody, 1:500, Cell Signaling, USA) and a fluorescent streptavidin (AMCA, anti-biotin, 1/1000, Vector laboratories) were applied overnight at 4°C. Slices were mounted with fluoromount and stored at 4°C in the dark. Pictures of POMC cells were captured using a confocal microscope with objective 63X (SP8-STED, Leica) setting fixed acquisition parameters. Sampling rate respected the optic resolution of the system (voxel size: 0.0819×0.0819×0.2985 μm^3^) and z-axis stacks were processed (around 20 images per cells). Images were acquired separately for each wavelength (blue=405nm and far red=647nm) and then merged for analysis. Using Imaris 8.1.2 software (Bitplane, Switzerland), POMC cell body volume (default settings for surface reconstruction) and individual GAD65/67 staining spots (0.4 μm diameters dots, threshold of 3500 pixels set taking into account background signal obtained from staining carried out in the absence of the primary antibody) were automatically reconstructed in 3D. A dedicated analysis module allowed making a clear distinction between the GAD65/67 spots located around the cell and the ones present within the cell body of the POMC neuron.

### 2.15. Statistics

Statistical analyses were performed using Prism, versions 6 and 8 (Graphpad, USA). All values, unless stated otherwise in the figure legends, are reported as means ± SEM. Data were analyzed by unpaired or paired Student’s t-tests, Mann-Whitney tests, one-way or two-way ANOVAs, as appropriate. Repeated measures for matched individuals were used when same animals underwent different treatments. Significant ANOVAs were followed by Fisher LSD post-hoc test. A Kruskal-Wallis test followed by Dunn’s post-test was used to assess differences in the distance to the ventricle across the different POMC subpopulations. To describe rostro-caudal changes in the distance to the ventricle for each subpopulation, linear regression analysis was performed using R followed by ANOVA. A Kruskal-Wallis test was also used for the analysis of the c-Fos expression in the different POMC neurons subpopulations in response to palatable food. Single-cell RNA-seq data for the various *Pomc*^+^ cells subgroups was analyzed using Student’s t-test on log-transformed mRNA copy data. P<0.05 denote statistical significance. Detailed statistical analysis is provided in Supplementary Tables S4 and S5.

## 3. Results

### 3.1. POMC neurons with specific neurotransmitter profiles have distinct spatial distribution and molecular fingerprints

POMC neurons are molecularly diverse (Campbell et al., 2017; Henry et al., 2015; Lam et al., 2017) and include different subtypes, also based on the neurotransmitter they produce (Dicken et al., 2012; Hentges et al., 2004; Hentges et al., 2009; Wittmann et al., 2013). Accordingly, triple fluorescent *in situ* hybridization (FISH) analysis in brains of adult C57BL/6J mice revealed different hypothalamic POMC subpopulations depending on whether or not POMC cells expressed the GABA synthesizing enzyme GAD67 and/or the glutamatergic marker vesicular glutamate transporter 2 (vGlut2, Figure 1A). Different POMC subpopulations also had specific spatial distribution, as assessed by analyzing their rostro-caudal distribution and the distance from the third ventricle (Figures 1B, 1C and S1A in Supplemental Material). POMC/glutamatergic neurons were generally furthest from the ventricle and had a rostro-caudal decrease, while POMC/GABAergic neurons were closest to the ventricle and had a rostro-caudal increase (Figures 1B, 1C and S1A). POMC GABA/glutamatergic cells and POMC cells negative for the 2 neurotransmitters were somewhat in between these two populations (Figures 1B, 1C and S1A). To then align POMC neurons to GABA, glutamate or mixed neurotransmitters expression, we analyzed publicly available single-cell RNA sequencing data of POMC neurons from the hypothalamus of C57BL/6N mice [GEO accession number: GSE74672, (Romanov et al., 2017)] by means of differential expression profiling (Figures 1D and S2). Of the 28 POMC cells examined, 7 exclusively expressed *Slc17a6*, coding for vGlut2; 11 exclusively contained *Gad1* and *Gad2*, coding for the GABAergic markers GAD67 and GAD65, respectively, and 10 were positive for both glutamatergic and GABAergic markers (Figure 1D). Two out of 11 POMC/GABAergic neurons were also positive for *Npy* only, and for *Npy* and *Agrp*, implying that *∼*18% of POMC/GABAergic neurons co-express these orexigenic neuropeptides. FISH analysis further confirmed co-localization between POMC and AgRP mRNA in the hypothalamus of adult mice, highlighting the co-expression of these two neuropeptides in the same cell (Figure S1B). POMC/GABAergic and POMC/glutamatergic subpopulations had non-overlapping gene expression profiles (Figures 1D and S2), while the mixed GABAergic/glutamatergic population expressed molecular markers typical of both POMC/GABAergic and POMC/glutamatergic cells (Figures 1D and S2). mTOR and associated proteins such as rptor and rictor (Saxton and Sabatini, 2017), as well as CB_1_R were detected in all POMC subpopulations (see GEO: GSE74672). Differential gene expression analysis done on POMC/GABAergic *vs.* POMC/glutamatergic subpopulations revealed that these two groups of neurons are substantially different in terms of the molecular machinery controlling substrates use and synaptic activity. POMC/GABAergic neurons were enriched in the expression of genes regulating glucose and lipid metabolism, protein transcription and catabolism (Table 1).Whilst POMC/glutamatergic neurons had increased expression of genes involved in synaptic vesicular function, purine metabolism and GTP signaling (Table 1). Thus, hypothalamic POMC neurons include different neurotransmitter-type subpopulations, with specific spatial positions and molecular signatures.

**Figure 1.**
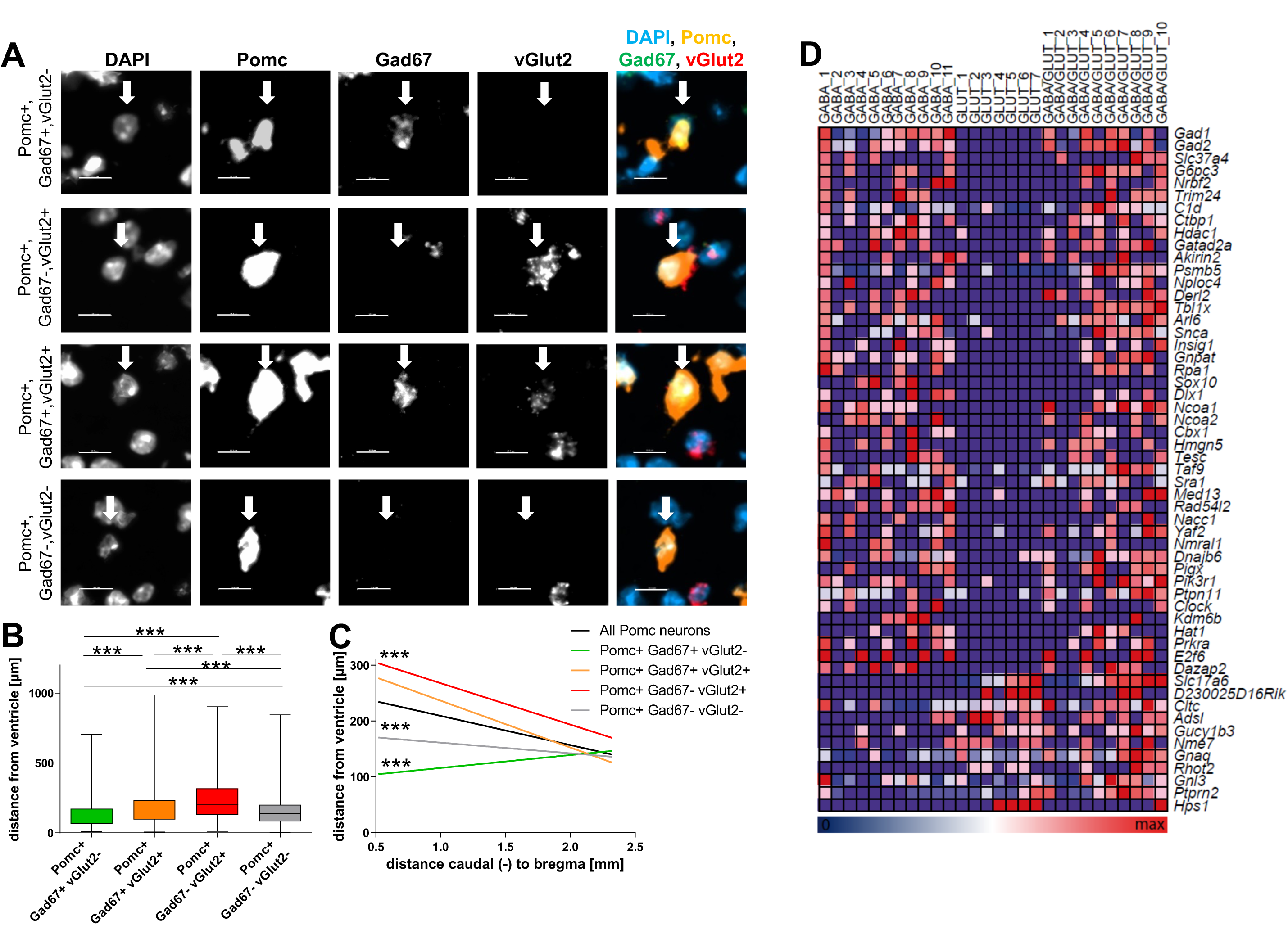
POMC neurons include GABAergic and glutamatergic subtypes with distinct spatial distribution and molecular fingerprints. (**A**) Representative images of triple FISH, showing different hypothalamic POMC cells expressing or not GABAergic (Gad67) and glutamatergic (vGlut2) markers. A total of 3914 cells in the ARC of 4 C57BL/6J mice were analyzed in B and C. Scale bar: 15µm. (**B**) Min to max plot showing that all subpopulations differed significantly concerning distance from the ventricle irrespective of rostro-caudal position (Kruskal-Wallis ANOVA followed by Dunn’s post-test). (**C**) Other than the GABA/glutamatergic POMC neurons, all subpopulations differed significantly from the regression line obtained for all analyzed POMC neurons (p values ≤ 4.6*10^-11^, ANOVA). (**D**) Neurotransmitter type-specific single-cell transcriptome profiling of hypothalamic *Pomc*^+^ neurons. The heat map illustrates the differential expression of several genes depending on whether *Pomc*^+^ cells are only GABAergic (11 cells, column 1 to 11 from the left), glutamatergic (7 cells, columns 12 to 18 from the left) or GABA/glutamatergic (10 cells, columns 19 to 28 from the left). The color scale bar shows the color coding of the single-cell expression level. Data are normalized to maximum single-cell level in the dataset for every gene. On the right: gene names. ***p<0.001. See also Table 1, Table S4 and Figures S1 and S2.

**Table 1.** Gene annotation enrichment analysis of differentially expressed genes in POMC/GABAergic and POMC/glutamatergic neurons.

|  | Database | Term | Fold Enrichment | Count | p-value |
| --- | --- | --- | --- | --- | --- |
| **Enriched in POMC/GABA Neurons** | Gene Ontology | Glutamate decarboxylase activity | 67.11 | 2 | 2,94E-02 |
|  | Gene Ontology | Glucose-6-phosphate/hexose phosphate transport | 45.07 | 2 | 4,35E-02 |
|  | Gene Ontology | Transcriptional repressor complex | 11.75 | 5 | 7,79E-04 |
|  | Gene Ontology | Proteasomal ubiquitin-dependent protein catabolic process | 9.01 | 4 | 9,52E-03 |
|  | Gene Ontology | Neutral lipid metabolic process | 6.01 | 4 | 2,84E-02 |
|  | Gene Ontology | Glycerol ether metabolic process | 6.01 | 4 | 2,84E-02 |
|  | Gene Ontology | Triglyceride metabolic process | 5.63 | 3 | 9,81E-02 |
|  | Gene Ontology | Chromatin binding | 4.33 | 10 | 5,03E-04 |
|  | Gene Ontology | Regulation of gene-specific transcription | 4.04 | 4 | 7,59E-02 |
|  | Gene Ontology | Transcription coactivator activity | 4.03 | 6 | 1,65E-02 |
|  | Gene Ontology | Transcription cofactor activity | 3.82 | 12 | 3,10E-04 |
|  | Gene Ontology | Transcription repressor activity | 3.73 | 10 | 1,45E-03 |
|  | Gene Ontology | Transcription corepressor activity | 3.73 | 4 | 9,12E-02 |
|  | Gene Ontology | Glycerolipid metabolic process | 3.14 | 6 | 4,19E-02 |
|  | Gene Ontology | Ubiquitin-dependent protein catabolic process | 2.88 | 6 | 5,74E-02 |
|  | Gene Ontology | Transcription factor binding | 2.80 | 12 | 3,82E-03 |
|  | Gene Ontology | Negative regulation of transcription | 2.54 | 14 | 3,56E-03 |
|  | Gene Ontology | Negative regulation of gene expression | 2.47 | 15 | 3,04E-03 |
|  | Gene Ontology | Transcription factor complex | 2.25 | 8 | 6,50E-02 |
| **Enriched in POMC/Glut Neurons** | Gene Ontology | Synaptic vesicle membrane | 34.64 | 2 | 5,48E-02 |
|  | Gene Ontology | Synaptic vesicle membrane | 34.64 | 2 | 5,48E-02 |
|  | Gene Ontology | Clathrin coated vesicle membrane | 27.42 | 3 | 5,04E-03 |
|  | Gene Ontology | Cytoplasmic vesicle membrane | 11.35 | 3 | 2,72E-02 |
|  | Kegg pathway | Purine metabolism | 7.31 | 3 | 5,46E-02 |
|  | Gene Ontology | GTP binding | 4.42 | 4 | 5,68E-02 |
|  | Gene Ontology | Vesicle | 3.17 | 5 | 6,59E-02 |

### 3.2. mTORC1 activity oppositely regulates POMC/GABAergic and POMC/glutamatergic neurons

Detection of energy availability by POMC neurons is essential for their role in the regulation of food intake (Cota et al., 2007; Krashes et al., 2016). Among the possible molecular mechanisms involved, mTORC1 activity is known to be a cellular readout of energy availability (Haissaguerre et al., 2014; Saxton and Sabatini, 2017). We therefore asked whether modulation of mTORC1 signaling might differentially regulate the activity of distinct subpopulations of POMC neurons, as identified through our neuroanatomical and transcriptomic analyses. Thus, we monitored POMC neuronal firing in hypothalamic slices from POMC-YFP mice under basal conditions and after bath application of the mTORC1 inhibitor rapamycin (RAPA, 200 nM), which mimics an acute cellular negative energy state (Haissaguerre et al., 2014; Saxton and Sabatini, 2017).

Taken all recorded cells together, RAPA had no effect on firing frequency (Figure S3A, n=117). However, individual firing analysis allowed identifying distinct POMC neuronal populations that responded differently to RAPA: (i) POMC neurons inhibited by RAPA (RAPA^inh^, Figures 2A, 2B, n=41/117, 35.1%), (ii) POMC neurons activated by RAPA (RAPA^act^, Figures 2A, 2C, n=37/117, 31.6%), and (iii) POMC neurons not responding to RAPA (RAPA^ns^, Figures 2A, 2D, n=39/117, 33.3%).

**Figure 2.**
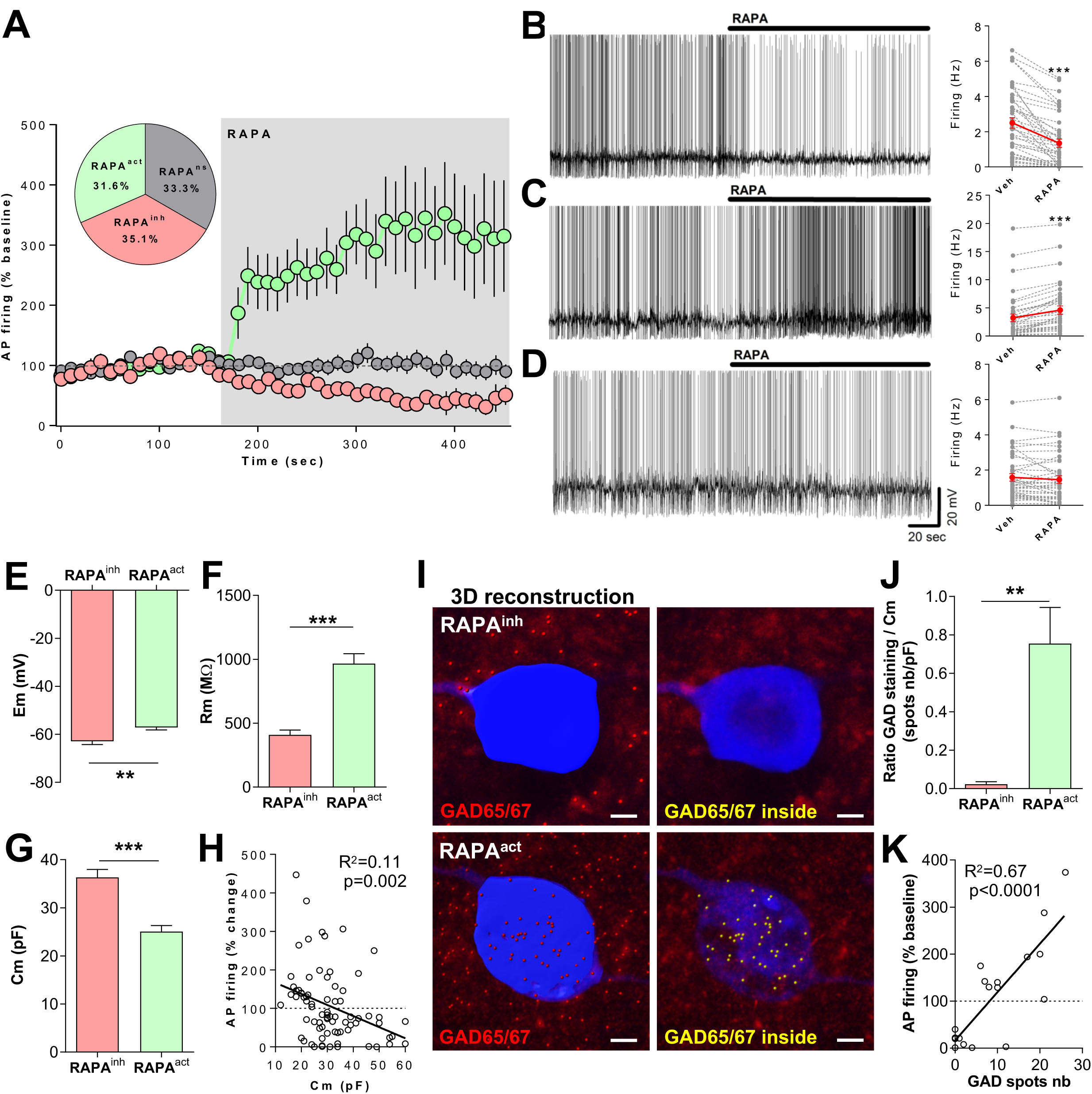
mTORC1 blockade oppositely regulates POMC neurons activity depending on neurotransmitter cell type. (**A**) Repartition of the tested POMC neurons as a function of their response to rapamycin (RAPA, 200nM). (**B**-**D**) Representative traces of POMC neurons firing during RAPA perfusion recorded in whole-cell current clamp configuration and corresponding firing changes in response to RAPA (right side). RAPA decreases (B, n=41/117 cells), increases (C, n=37/117 cells) or has no effect (D, n=39/117 cells) on POMC firing (46 mice analyzed in total). (**E**-**G**) RAPA^act^ POMC neurons (n=37) have increased membrane resting potential (Em; E), and membrane resistance (Rm; F), and decreased cell capacitance (Cm; G) as compared to RAPA^inh^ POMC neurons (n=41). (**H**) Change in cell firing in response to RAPA is correlated with Cm (n=78 cells). (**I**) Representative 3D-reconstruction from Z-stack confocal images of RAPA^inh^ and RAPA^act^ POMC neurons used for electrophysiological recordings. Reconstruction of cell body (biocytin: blue surface) and GAD staining (GAD65/67: red spots, left panel), and corresponding images showing only the GAD spots present inside the cell body surface (yellow, right panel). (**J**) Number of GAD65/67 spots inside the cell surface over membrane capacitance is higher in RAPA^act^ POMC neurons as compared to RAPA^inh^ POMC neurons (n=7-11 cells). (**K**) Change in action potential firing under RAPA correlates with the number of GAD spots present inside the cell (n=18 cells). Data analyzed by paired t-test (B-D), unpaired t-test (E-G, J) and linear regression (H, K).**p<0.01; ***p<0.001. Scale bar in I: 3 μm. See also Figure S3 and Table S4.

These subpopulations had specific electrophysiological properties, allowing prediction of their functional properties. In particular, RAPA^act^ POMC neurons had higher membrane resting potential (Em) and resistance (Figures 2E, 2F), and lower membrane capacitance (Cm, Figure 2G) than RAPA^inh^ POMC cells. RAPA^ns^ cells had characteristics that were in-between RAPA^act^ and RAPA^inh^ POMC neurons (data not shown). RAPA depolarized RAPA^act^ POMC cells while hyperpolarizing RAPA^inh^ POMC cells (Figure S3B). An inverse correlation was found between the recorded cells’ Cm and the firing changes induced by RAPA (Figure 2H).

Cm is an indicator of cell size (Hentges et al., 2009) and relatively smaller size seems the most reliable marker to distinguish POMC/GABAergic neurons *vs.* POMC/glutamatergic ones (Hentges et al., 2004; Hentges et al., 2009). Based on the Cm, we hypothesized that RAPA^inh^ POMC cells were likely glutamatergic, while RAPA^act^ POMC neurons were GABAergic. To confirm this, recorded neurons were filled with biocytin and subsequently labelled with an antiserum directed against the GABAergic markers GAD65/67. RAPA^act^ POMC neurons had significant GAD65/67 staining, whereas RAPA^inh^ POMC neurons did not overcome background staining levels (Figures 2I, 2J, S3C and S3D). GAD65/67 staining inside the cell positively correlated with the firing changes induced by RAPA (Figure 2K) and was inversely associated with the Cm of the analyzed cells (Figure S3E). A negative correlation was also found between the firing changes caused by RAPA and the Cm (Figure S3F), overall implying that small POMC cells activated by RAPA are likely GABAergic.

Then, by using optogenetics, we evaluated whether the acute inhibition of mTORC1 activity could alter POMC-driven neurotransmission. Among the different brain structures receiving POMC neurons projections, we studied this effect on parvocellular neurons of the PVN, since they are one of the targets of POMC neurons for the control of food intake (Singru et al., 2012; Mazier et al., 2019).

Using the FLEX system (Atasoy et al., 2008) in POMC-Cre neurons (Balthasar et al., 2004), we expressed the photoactivable channelrhodopsin-2 (ChR2) (Atasoy et al., 2008) and recorded light-evoked ARC POMC synaptic inputs onto PVN parvocellular neurons (Figure 3A), which were recognized based on their electrophysiological properties (Luther and Tasker, 2000). About 1 in 10 parvocellular neurons were synaptically connected to terminals arising from infected POMC neurons (31/321 cells; 9.66%). Pharmacological assays on light-evoked synaptic transmission were performed in 17 parvocellular cells (Figures 3B, S4 and Table S2). The majority of the POMC terminals connecting onto parvocellular neurons released pure glutamatergic inputs (58.82%; 10/17), whereas the remaining recorded cells received pure GABAergic (17.65%; 3/17) or mixed GABA/glutamate inputs (23.53%; 4/17) (Figure 3B). To test the involvement of mTORC1 in the control of neurotransmitter release from POMC terminals onto parvocellular neurons, we challenged light-evoked neurotransmission by adding RAPA to the perfusate. Acute inhibition of mTORC1 activity reduced light-evoked excitatory post-synaptic currents (eEPSCs) amplitude (Figures 3C, 3D), while increasing light-evoked inhibitory post-synaptic currents (eIPSCs) amplitude (Figures 3C, 3E). Thus, mTORC1 activity oppositely regulates POMC/GABAergic and POMC/glutamatergic transmission.

**Figure 3.**
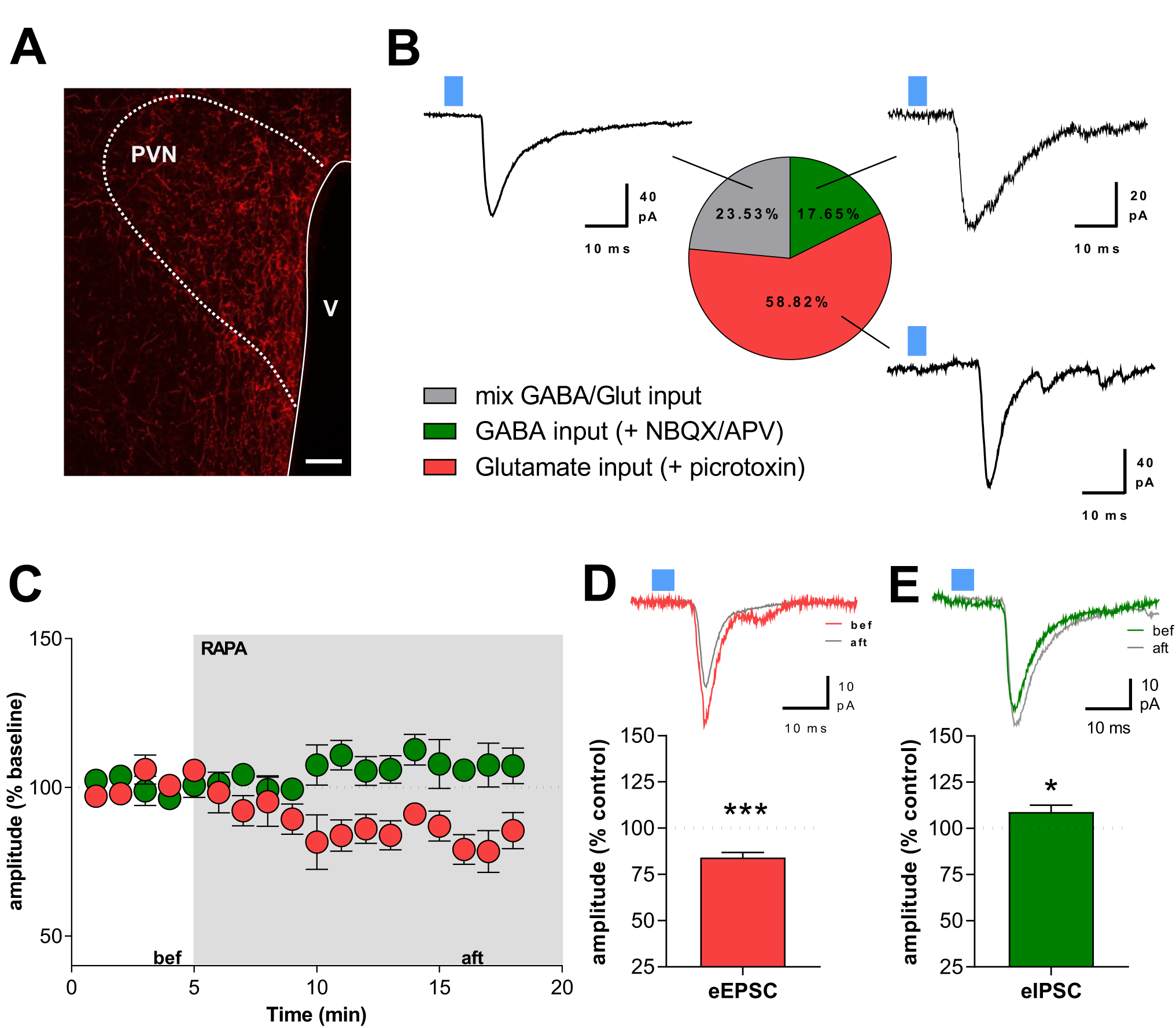
mTORC1 activity modulates POMC neurons neurotransmission onto PVN parvocellular cells. (**A**) Representative image of mCherry-positive, ChR2-expressing projections from ARC POMC neurons onto the hypothalamic paraventricular nucleus (PVN). (**B**) Distribution of the neurotransmission type recorded in PVN parvocellular neurons after photostimulation (n=17 cells) and representative traces of light-evoked post-synaptic currents for each neurotransmission type. POMC neurons mainly release glutamate (58.82%, 10/17 cells) onto parvocellular neurons; pure GABAergic (17.65%; 3/17 cells) and mixed GABA/Glutamate (23.53%, 4/17 cells) connections were also found. (**C**) Amplitude time course of light-evoked EPSC (red; n=6 cells) and IPSC (green; n=5 cells) normalized to baseline. RAPA decreases eEPSCs amplitude (**D**), and increases eIPSCs amplitude (**E**). Representative traces of eEPSC (D; top) and eIPSC (E; top) before (bef, in C) and after (aft, in C) RAPA application. Data in D and E analyzed by paired t-test. *p<0.05, ***p<0.001. V: 3^rd^ ventricle. Scale bar in A: 50 μm. Blue square: light. See also Figure S4 and Tables S2 and S4.

### 3.3. mTORC1 signaling in POMC neurons regulates food intake

Hypothalamic POMC neurons are typically found activated (increased c-Fos expression) up to 2h after refeeding, a condition engaging *α*-MSH dependent MC4R signaling in the PVN, which then helps terminate the meal (Balthasar et al., 2005; Fekete et al., 2012; Singru et al., 2007). To test the relevance of the mTORC1 pathway in modulating POMC neurons-associated feeding responses, we first evaluated the impact of an intracerebroventricular (icv) administration of RAPA on refeeding in C57BL/6J mice. RAPA, which was administered just before access to food, acutely increased food intake, particularly at 2h (Figure 4A). This change in food intake was associated with changes in c-Fos expression in POMC neurons, which were activated by refeeding, in agreement with (Fekete et al., 2012; Singru et al., 2007) and partly inhibited by icv RAPA administration, as compared to fasting (Figure 4B,4D). Changes in mTORC1 activity followed similar trends, with an increased phosphorylation of mTORC1 downstream effector S6 ribosomal protein (p-S6) (Saxton and Sabatini, 2017), in activated POMC neurons in refeeding, which was blunted by RAPA (Figure 4C-D). This evidence therefore suggests that inhibition of mTORC1 activity in POMC neurons may be relevant for the effect of RAPA on food intake.

**Figure 4.**
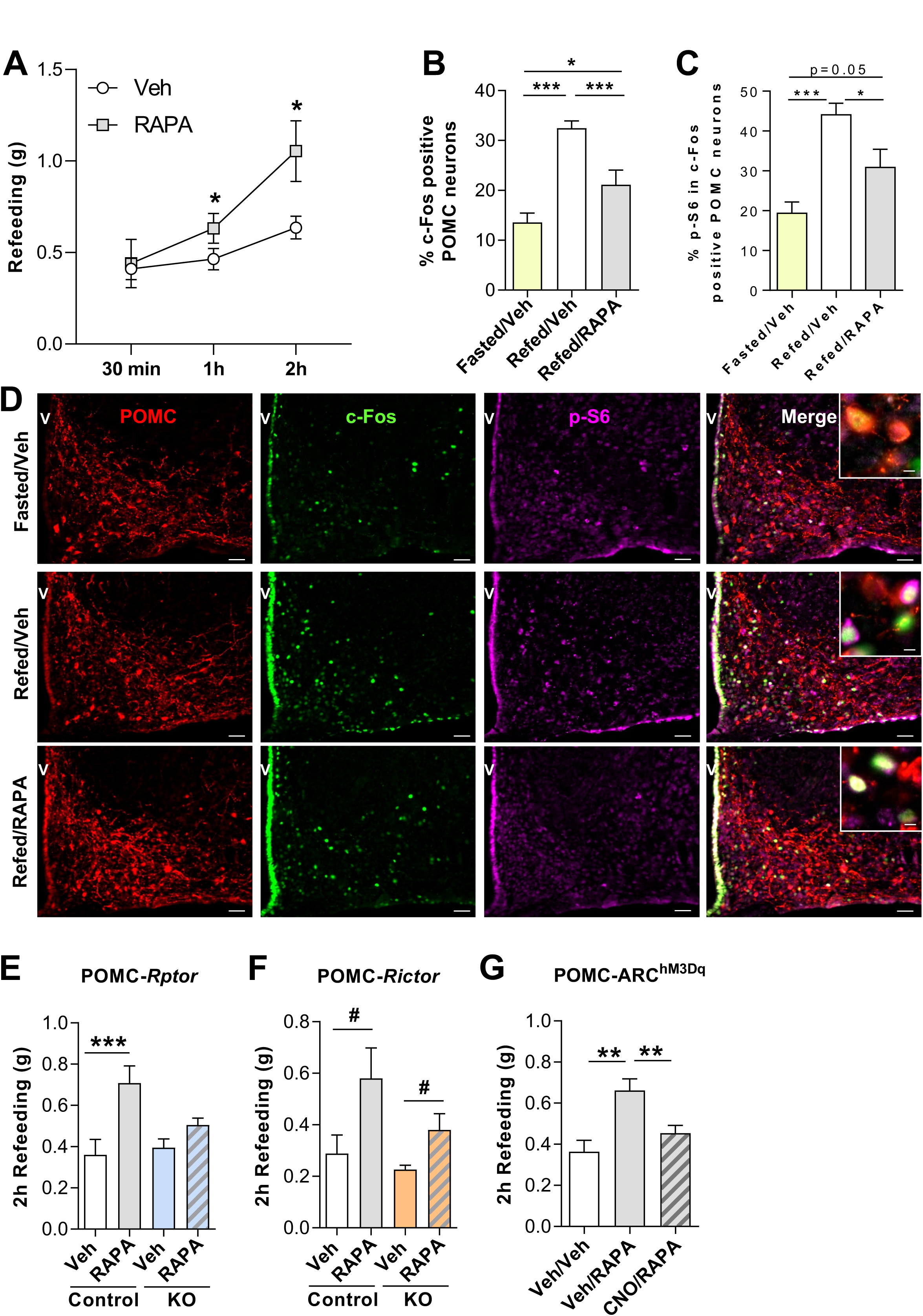
mTORC1 in POMC neurons controls food intake. Refeeding (fasting-induced food intake) of C57BL/6J mice treated icv with RAPA or its vehicle (**A**, n=5 mice per group). Quantification of c-Fos (**B**) and of p-S6 (**C**) labelling in POMC-positive neurons in the ARC and related representative images (**D**) obtained from C57BL/6J mice fasted for 24h or refed for 90 min and treated icv with RAPA or its vehicle (n=4-5 mice per group). 2h Refeeding of POMC-*Rptor*-Control and KO littermates (**E**, n=6-9 mice per group), and POMC-*Rictor*-Control and KO littermates (**F**, n=5 mice per group) treated icv with RAPA or its vehicle. (**G**) Acute ip CNO injection inhibits icv RAPA-induced 2h hyperphagia in POMC-ARC^hM3Dq^ mice (n=5 mice per group). Data analyzed by RM two-way ANOVA (A,E, F), one-way ANOVA (B, C) or RM one-way ANOVA (G) followed by Fisher LSD post-hoc test.*p<0.05; **p<0.01; ***p<0.001. ^#^p<0.05, treatment effect. Scale bar in D: 50µm, and 10µm for smaller inset. V: 3rd ventricle. See also Figure S5 and Table S4.

To test this hypothesis, we probed the effect of the drug in POMC-*Rptor*-KO mice, which carry a specific deletion of the gene *rptor*, necessary for functional mTORC1 signaling (Hara et al., 2002), in POMC neurons (Haissaguerre et al., 2018). Of note, these mice do not show any alteration in their basal, unstimulated, feeding behavior (Haissaguerre et al., 2018). Acute RAPA icv administration caused hyperphagia in POMC-*Rptor*-controls, but not in KO littermates (Figure 4E). In contrast, icv administration of RAPA to mutant mice lacking the gene *rictor* [required for mTORC2 formation (Sarbassov et al., 2004)] in POMC neurons (POMC-*Rictor*-KO mice; Figures S5A, S5B) still increased food intake (Figure 4F). This genetic model also had unstimulated food intake similar to their control littermates (Figure S5C). Thus, acute pharmacological blockade of mTORC1 activity causes hyperphagia by targeting mTORC1 in POMC neurons.

To further test whether POMC neurons activity may play a necessary role for RAPA-induced hyperphagia, we used the designer-receptors-exclusively-activated-by-designer-drugs (DREADD) approach (Alexander et al., 2009) in order to explore the effects of the mTORC1 inhibitor in response to the activation of POMC neurons. The gene encoding the evolved human M3-muscarinic receptor fused to the fluorescent protein mCherry (hM3Dq-mCherry) was selectively expressed in POMC neurons by injecting a Cre-inducible adeno-associated viral vector (AAV-DIO-hM3Dq-mCherry) into the ARC of POMC-Cre mutants to obtain POMC-ARChM3Dq mice [Figure S5D, (Zhan et al., 2013)]. As expected, systemic administration of the hM3Dq ligand clozapine-N-oxide (CNO, 1 mg/kg) increased the expression of the neuronal activity marker c-Fos in mCherry-positive POMC neurons (Figures S5E, S5F). c-Fos positive cells were also largely co-labeled by an antiserum detecting the mTORC1 downstream target p-S6 (Figures S5E, S5G). In agreement with (Koch et al., 2015; Zhan et al., 2013), systemic CNO administration did not acutely alter the food intake of POMC-ARC^hM3Dq^ mice (Figure S5H), but it fully prevented the hyperphagic effect of RAPA (Figure 4G). CNO was unable to block RAPA-induced hyperphagia in POMC-ARC^hM3Dq^-control mice, obtained by injecting AAV-DIO-hM3Dq-mCherry into the ARC of mice not expressing Cre in POMC neurons (Figure S5I).

Hence, acute inhibition of mTORC1 signaling increases food intake by likely altering the function of POMC neurons, which led us to investigate whether changes in POMC-derived neuropeptides and/or neurotransmitters could be involved in this phenomenon.

### 3.4. Acute inhibition of mTORC1 activity causes fasting-like changes in hypothalamic α-MSH and endocannabinoids levels

The POMC protein must be cleaved to generate active peptides that modulate food intake [Figure 5A and (Wardlaw, 2011)]. We measured hypothalamic content of POMC, its processing intermediates and cleavage products in 24-h fasted and 2-h refed mice treated icv with RAPA or its vehicle. Neither POMC nor its initial product adrenocorticotrophic hormone (ACTH) were altered by RAPA or refeeding (Table S3). However, hypothalamic levels of *α*-MSH were increased after refeeding relative to fasting, and RAPA administration just before access to food completely prevented this increase (Figure 5B). As mentioned earlier, *α*-MSH is implicated in determining satiety (Balthasar et al., 2005; Singru et al., 2007; Singru et al., 2012), while *β*-EP stimulates feeding (Koch et al., 2015). Also *β*-EP showed changes, albeit non-significant, similar to *α*-MSH in response to refeeding and to RAPA (Table S3). When using these data to calculate ratios of *α*-MSH/POMC, *α*-MSH/ACTH and *α*-MSH/*β*-EP, a significant relative reduction in hypothalamic *α*-MSH was found in refed, RAPA-treated mice as compared to vehicle (Figure 5C-E). Thus, acute inhibition of mTORC1 activity prevents the increase in hypothalamic *α*-MSH levels classically observed with food consumption (Krashes et al., 2016). Differently from *α*-MSH, hypothalamic endocannabinoids levels are high in fasting and low in refeeding (Kirkham et al., 2002) and CB_1_R activation counteracts behavioral and neuromodulatory effects of *α*-MSH (Mazier et al., 2019; Monge-Roffarello et al., 2014; Verty et al., 2004). We therefore reasoned that changes in hypothalamic *α*-MSH levels induced by the acute blockade of mTORC1 activity could be accompanied by opposite changes in hypothalamic endocannabinoid levels, which could have participated to the observed hyperphagia. As compared to fasting, refeeding dampened hypothalamic levels of the endocannabinoid anandamide (AEA) (Figure 5F). However, icv administration of RAPA just before access to food maintained hypothalamic AEA levels high, like in fasting (Figure 5F). RAPA had no effect on the hypothalamic (Figure 5G) or extra-hypothalamic (Table S3) content of the other endocannabinoid 2-arachidonoyl glycerol (2-AG) or on extra-hypothalamic levels of AEA (Table S3), independently of the feeding state. To determine whether RAPA-induced changes in hypothalamic *α*-MSH and AEA levels were causally linked, we co-administered RAPA icv together with the stable *α*-MSH analog MTII, used at a dose that did not have any effect on food intake or on hypothalamic AEA levels (0.02 μg icv, Figures S6A, S6B). In refed mice, MTII blocked the effect of RAPA on hypothalamic AEA levels (Figure 5H). Similarly, chemogenetic activation of POMC neurons prevented the effect of RAPA on hypothalamic AEA levels (Figure 5I). Hence, acute mTORC1 inhibition decreases hypothalamic *α*-MSH levels, which in turn leads to a hypothalamic increase in the endocannabinoid AEA.

**Figure 5.**
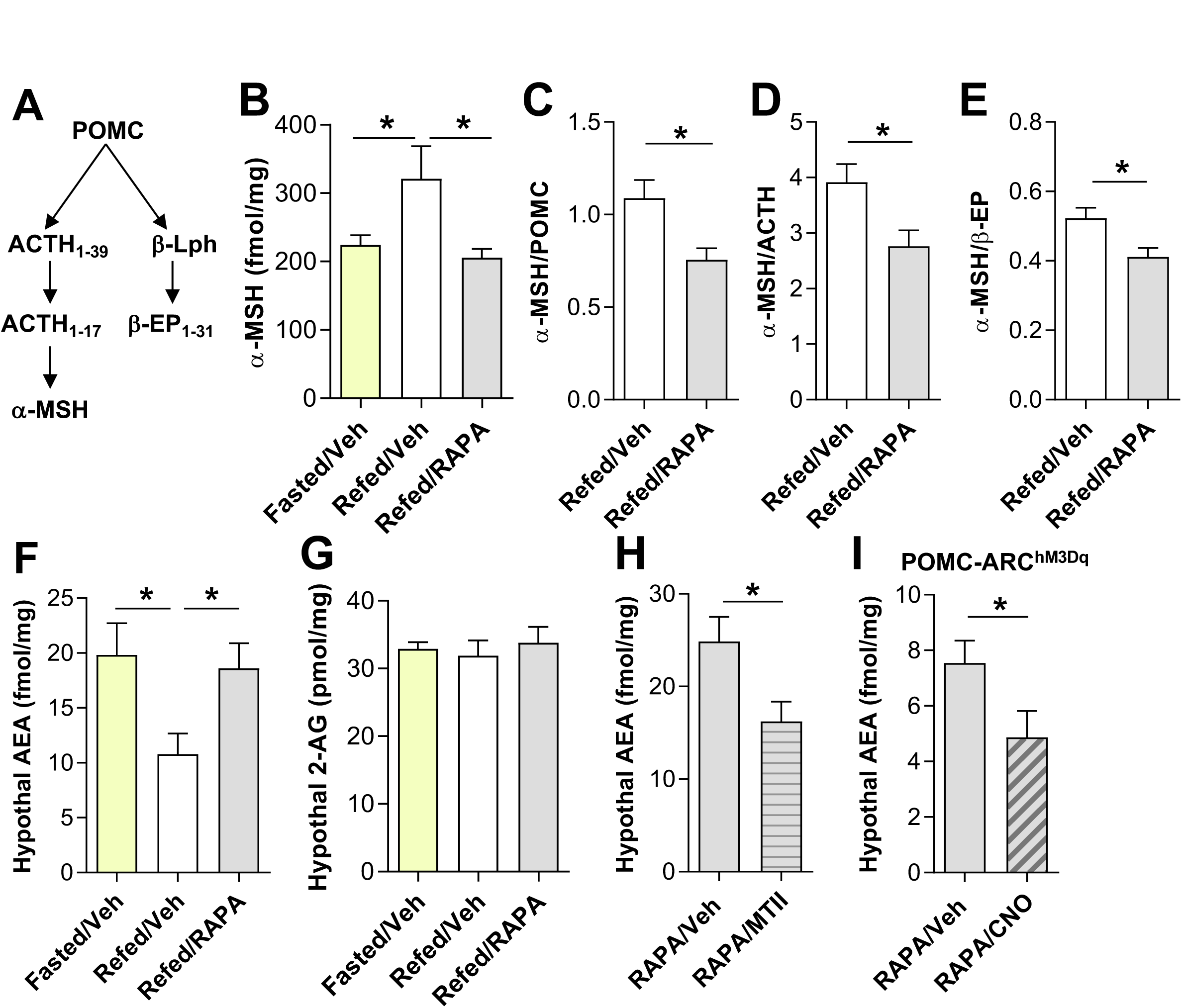
mTORC1 blockade induces fasting-like *α*-MSH and AEA changes in hypothalamus. (**A**) Schematic diagram illustrating hypothalamic POMC processing. (**B**) *α*-MSH content in the hypothalami of fasted or refed C57BL/6J mice treated icv with RAPA or its vehicle (n=8 mice per group). (**C**-**E**) Hypothalamic *α*-MSH/POMC (C), *α*-MSH/ACTH (D) and *α*-MSH/β-EP (E) ratios in 2-h refed C57BL/6J mice treated icv with RAPA or its vehicle (n=8 mice per group). (**F**, **G**) Effect of icv administration of RAPA or its vehicle on hypothalamic AEA (F) and 2-AG (G) content in fasted or refed C57BL/6J mice (n=3-5 mice per group). (**H**) Effect of the combined icv administration of the *α*-MSH analog MTII (0.02 µg) and RAPA on hypothalamic AEA content in refed C57BL/6J mice (n=6 mice per group). (**I**) Effect of the combined administration of CNO and RAPA on hypothalamic AEA content in refed POMC-ARC^hM3Dq^ mice (n=8-10 mice per group). Data analyzed by one-way ANOVA followed by Fisher LSD post-hoc test (B, F, G) or unpaired t-test (C-E, H, I) *p<0.05; **p<0.01; ***p<0.001. See also Figure S6 and Tables S3 and S4.

### 3.5. mTORC1-dependent recruitment of CB_1_R signaling restrains GABA release from POMC neurons

Endocannabinoids modulate food intake by acting onto CB_1_R (Mazier et al., 2015). To verify whether the hyperphagia induced by RAPA was due to increased endocannabinoid-CB_1_R signaling at the level of POMC neurons, we evaluated the effect of the mTORC1 inhibitor in mice lacking the expression of *CB_1_* in POMC neurons (POMC-*CB_1_*-KO mice) (Mazier et al., 2019). Deletion of *CB_1_* from POMC neurons did not alter *per se* 24h food intake (Figure S7A), or the refeeding response (Figure 6A). As expected, icv RAPA administration just before access to food caused hyperphagia in POMC-*CB_1_*-control mice (Figure 6A). Surprisingly, however, POMC-*CB_1_*-KO mice displayed greater hyperphagia than their controls in response to RAPA (Figure 6A), implying that CB_1_R on POMC neurons might actually inhibit food intake in response to acute changes in mTORC1 activity. Our past work has demonstrated that while activation of CB_1_R on glutamatergic neurons increases food intake, its activation on GABAergic terminals decreases it (Bellocchio et al., 2010). As RAPA stimulates the firing of POMC/GABAergic cells (Figures 2, 3), we hypothesized that deletion of CB_1_R in POMC neurons would not restrain anymore GABA release from POMC synaptic terminals, thereby further increasing food intake under acute mTORC1 blockade. Confirming this idea, co-administration of a sub-effective dose (0.03 μg/μl icv, Figures S7B, S7C)] of the GABAA receptor antagonist picrotoxin (Ptx), blocked the excessive RAPA-induced hyperphagia in POMC-*CB_1_*-KO mice, reducing their response to similar levels as RAPA-treated, POMC-*CB_1_*-control littermates (Figure 6A). Then, to confirm that lack of CB_1_R on POMC neurons alters POMC/GABAergic transmission, we evaluated miniature inhibitory post-synaptic currents (mIPSCs) frequency onto PVN parvocellular neurons. In agreement with the inhibitory role of CB_1_R activation on neurotransmitter release (Busquets-Garcia et al., 2018; Castillo et al., 2012), bath application of the CB_1_R agonist WIN 55,212-2 inhibited mIPSCs frequency in control littermates, but not in POMC-*CB_1_*-KO mice (Figure S7 C-E).

**Figure 6.**
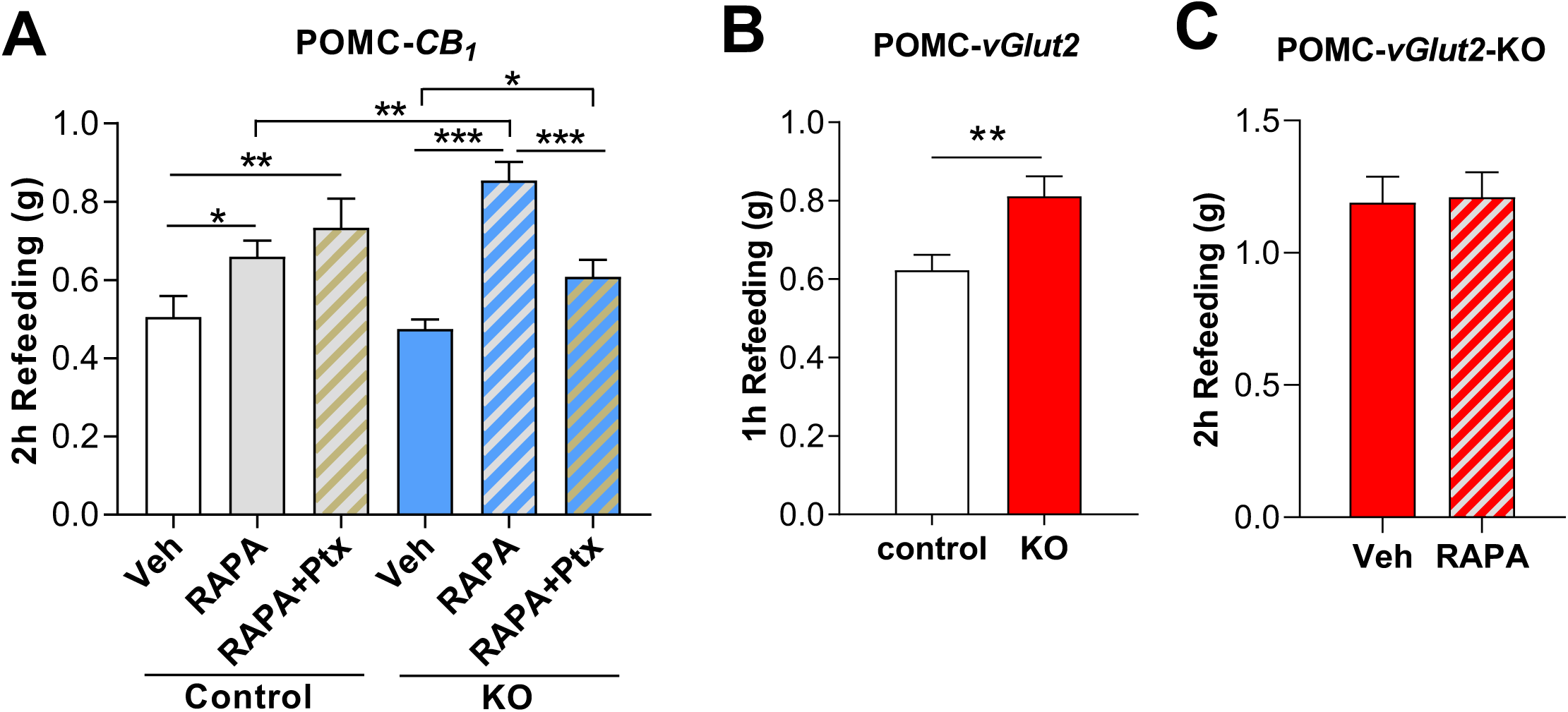
POMC/GABAergic and POMC/glutamatergic neurons oppositely regulate food intake. (**A**) Refeeding in POMC-*CB_1_*-Controls and KO littermates treated icv with RAPA, its vehicle or a combination of RAPA and the GABAA receptor antagonist picrotoxin (Ptx) (n=8-18 mice per group, two-way ANOVA followed by Fisher LSD post-hoc test, 2 experiments combined together). (**B**) Refeeding response in POMC-*vGlut2*-Control and KO littermates (n=8-14 mice per group, unpaired t-test) and (**C**) effect of an icv administration of RAPA or its vehicle on refeeding in POMC-*vGlut2*-KO mice (n=8 mice per group, paired t-test). *p<0.05; **p<0.01; ***p<0.001. See also Figures S7-S9 and Table S4.

Altogether, these data indicate that acute mTORC1 blockade activates POMC/GABAergic transmission, which drives feeding under the negative control of CB_1_R signaling. Hence, to further explore whether the POMC/GABAergic neuronal population may be involved in promoting feeding, we evaluated the activity of the different POMC neurons subpopulations in free-fed C57BL/6J mice exposed for 2h to palatable food. This experimental paradigm strongly increased food intake (Figure S8A), leading to increased c-Fos expression in POMC neurons (Figure S8B, S8C), and a high prevalence of c-Fos specifically in the POMC/GABAergic population, as compared to all other POMC neurons subpopulations (Figure S8C, S8D).

### 3.6. Inhibition of glutamate release from POMC neurons causes hyperphagia

Our electrophysiology studies also indicated that acute blockade of mTORC1 activity inhibited POMC/glutamatergic transmission. To finally evaluate *in vivo* the relevance of this effect for feeding behavior, we generated tamoxifen-inducible POMC-CreER^T2^/vGlut2^fl/fl^ mice (thereafter called POMC-*vGlut2*-KO). We used POMC-CreER^T2^-Ai6 reporter mice to confirm that tamoxifen induced recombination in POMC neurons (Figure S9A) and verified deletion of vGlut2 in hypothalamic samples of POMC-*vGlut2*-KO and control littermates by PCR (Figure S9B). As compared to their controls, POMC-*vGlut2*-KO mice had similar 24h food intake under unstimulated conditions (Figure S9C), but were hyperphagic in refeeding (Figure 6B), which was possibly also the result of the POMC/GABAergic component being still functional in this mouse model. Besides, POMC-*vGlut2*-KO mice did not respond to the icv administration of RAPA (Figure 6C). Thus, the genetic inhibition of glutamate signaling from POMC neurons recapitulates the hyperphagic phenotype caused by the acute blockade of mTORC1.

Overall, the data indicate that POMC/GABAergic and POMC/glutamatergic neurons counterbalance each other in an mTORC1-dependent fashion to modulate feeding.

## 4. Discussion

Over the past 20 years, several studies have outlined the functions of AgRP and POMC neurons in the context of energy balance, leading to the widespread notion that AgRP neurons increase food intake, while POMC neurons decrease it. The present data now indicate that the functional purpose of two distinct POMC neuronal subtypes, one GABAergic, the other glutamatergic, is to oppositely regulate food intake in response to changes in energy availability.

Indeed, our findings suggest that acute inhibition of mTORC1 activity, which artificially mimics low cellular energy levels, induces simultaneous activation of POMC/GABAergic neurons and inhibition of POMC/glutamatergic cells and melanocortin signaling. This leads to the recruitment of endocannabinoid-CB_1_R signaling, which likely hinders excessive GABA release from POMC neurons, overall resulting in hyperphagia. Based on this evidence, it would be anticipated that increased mTORC1 activity in POMC neurons as a result of higher energy availability due to food intake (Cota et al., 2006; Dagon et al., 2012; Haissaguerre et al., 2018), would help terminate the meal through a chain of events, including inhibition of POMC/GABAergic and endocannabinoid-CB_1_R activity, and stimulation of *α*-MSH and POMC/glutamatergic signaling. Accordingly, deletion of glutamatergic signaling from POMC neurons of adult mice behaviorally recapitulates the effects obtained with the pharmacological blockade of mTORC1.

Thus, the evidence we provide uncovers a more complex and dynamic role for POMC neurons in the control of food intake than has been appreciated so far, suggesting that the dichotomic view of AgRP *vs.* POMC neurons in the control of food intake is one piece of a far more complex picture. Fiber photometry studies have recently shown that the sensory detection of food is sufficient to immediately switch off AgRP neurons and activate POMC neurons (Chen et al., 2015). However, these studies did not address the neurotransmitter nature of the POMC cells involved. Our data now support the hypothesis that POMC neurons, thanks to their heterogeneity, may sequentially regulate feeding by initially favoring ingestion of food via POMC/GABAergic cells and then help terminate the meal through the POMC/glutamatergic component.

Our neuroanatomical analyses show that distinct POMC neurotransmitter cells subtypes have specific spatial location in the ARC. This is consistent with recent studies describing distinct spatial distribution of POMC neurons based on the expression of GABAergic and glutamatergic markers (Jones et al., 2019). Spatial location could be eventually associated with specific projections targets, as it has been demonstrated for AgRP neurons (Betley et al., 2013). Transcriptomic profiling in the hypothalamus of C57BL/6 mice also proves that POMC/glutamatergic and POMC/GABAergic subpopulations have distinct neurochemical features, as POMC/GABAergic cells have higher glycolytic capacity and expression of transcription factors and lower expression of synaptic proteins relative to POMC/glutamatergic neurons. We found that a small fraction of POMC/GABAergic neurons also co-express *NPY* and/or *AgRP*, implying that there is a subgroup of hypothalamic GABAergic cells that contains both POMC and AgRP in adulthood. This observation agrees with previous transcriptomic data obtained using reporter mice to trace POMC and AgRP neurons (Lam et al., 2017), and likely reflects the shared developmental origins of POMC and NPY/AgRP cells (Padilla et al., 2010), indicating that these features are present in C57BL/6 mice and are somehow conserved during adult life. Such a ‘mixed’ phenotype of POMC neurons also represents a limitation of our study, since the genetic models we used, generated with a POMC-Cre transgene expressed during either development or adulthood, may leave out the possibility that some of the effects observed are due to POMC-derived non-POMC GABAergic cells, namely NPY/AgRP cells.

Our study also reveals that there is a POMC/GABAergic component involved in the stimulation of food intake, as pointed out by our investigations in POMC-*CB_1_*-KO mice and the observed association between consumption of palatable food and activation of POMC/GABAergic neurons in C57BL/6 mice. However, further studies are needed in order to pinpoint the exact role of the POMC/GABAergic population in the regulation of food intake and more specifically of POMC-NPY/AgRP cells in this context.

Besides, it has been proposed that the proportion of POMC/glutamatergic and POMC/GABAergic neurons varies with age (Dennison et al., 2016). In this regard, future longitudinal studies will address the interesting possibility that the dual GABA/glutamatergic population that we have also identified might be indicative of a switch in neurotransmitter phenotype over time and/or it represents another way for POMC neurons to fine-tune their actions on target circuits. Noteworthy, the heterogeneity in neurotransmitter type might explain the inability to observe acute, rapid changes in food intake reported in several studies using opto- or chemo-genetics to modulate POMC neurons activity [present findings and (Aponte et al., 2011; Koch et al., 2015; Zhan et al., 2013)]. Likewise, the fact that POMC-*Rptor*-KO, POMC-*Rictor*-KO and POMC-*CB_1_*-KO mice did not have *per se* a phenotype during refeeding could possibly be due to the expression of the deleted genes across the different POMC neuronal subpopulations and the counterbalancing physiological functions of these subpopulations. Alternatively, the lack of phenotype could be due to compensation. This remains a possibility, albeit specific and different effects on food intake were observed in these models in response to pharmacology.

Our optogenetic electrophysiology studies also demonstrate that POMC/glutamatergic and POMC/GABAergic neurons functionally project to parvocellular neurons of the PVN, contributing to the activity of this brain region. This evidence agrees with previous studies illustrating both synaptic GABA and glutamate release from POMC neurons (Atasoy et al., 2008; Hentges et al., 2004; Dicken et al., 2012; Mazier et al., 2019). However, another study in which optogenetics was used to assess POMC/glutamatergic transmission onto the PVN was unable to find any, rather showing that such transmission depends upon ARC vGlut2-positive neurons, 44% of which were nevertheless POMC neurons (Fenselau et al., 2017). These differences could be due to the genetic models used, the interval between the virally-mediated delivery of the opsin and actual electrophysiological recordings, and the number and type of PVN cells analyzed. For instance, Fenselau et al. did not evaluate transmission onto parvocellular neurons, but studied all MC4R neurons (Fenselau et al., 2017). Besides, our study demonstrates that POMC/glutamatergic activity is particularly relevant to inhibit food intake. Accordingly, others have shown that mice with constitutive deletion of vGlut2 in POMC neurons are prone to develop diet-induced obesity (Dennison et al., 2016).

After the discovery of the involvement of the mTORC1 pathway in the hypothalamic regulation of energy balance (Cota et al., 2006), several studies have investigated the role of this pathway in AgRP and POMC neurons (Burke et al., 2017; Caron et al., 2016; Mori et al., 2009; Smith et al., 2015; Yang et al., 2012). The current findings now demonstrate that mTORC1 is an important molecular switch controlling recruitment of POMC neurons in response to energy availability and that, depending on the neurotransmitter type, mTORC1 activity leads to opposite effects on neurotransmission and food intake. Previous investigations have shown that the mTOR pathway has a key role in neurotransmission, particularly in the hippocampus, by modulating dendrite length and spine density, and by modifying the function and coding properties of individual synapses (Weston et al., 2012). While further studies will have to investigate the molecular mechanisms underlying the effects of rapamycin on POMC neuronal activity and transmission, it is worth mentioning that mTOR activity differentially regulate glutamatergic and GABAergic transmission in other brain areas (Weston et al., 2012).

Changes in POMC neuronal activity induced by mTORC1 are then associated with changes in hypothalamic levels of *α*-MSH and the endocannabinoid AEA, which eventually modulate neurotransmitter release and the feeding response. Thus, mTORC1 activity coordinates neuropeptidergic-(i.e. levels of *α*-MSH) and neurotransmitter-dependent responses of POMC neurons in relation to food intake. In agreement with previous evidence (Mazier et al., 2019; Monge-Roffarello et al., 2014; Verty et al., 2004), the recruitment of melanocortin signaling is upstream endocannabinoid-dependent CB_1_R activation. The latter not only inhibits POMC/glutamatergic transmission (Mazier et al., 2019), but it also hinders the POMC/GABAergic neuronal subpopulation, as suggested by our studies combining the in vivo administration of the GABAA antagonist picrotoxin in POMC-*CB_1_*-KO mice. The phenomena described can be related to the classic effects of presynaptic CB_1_R and are consistent with our past work demonstrating opposite roles of CB_1_R signaling on food intake depending on the neurotransmitter cell type (Bellocchio et al., 2010; Soria-Gomez et al., 2014b). Although not investigated in the current study, it is likely that the potential endocannabinoid-dependent retrograde suppression of GABA release and consequent modulation of food intake may involve POMC/GABAergic projections onto the PVN, since other studies have demonstrated that blockade of CB_1_R signaling in the PVN stimulates food intake during refeeding (Soria-Gomez et al., 2014a). Recent work has also shown that POMC neurons drive the hyperphagia induced by exogenous cannabinoids in sated mice through a mechanism that relies on CB_1_R modulation of presynaptic inputs onto POMC cells and on CB_1_R-dependent mitochondrial adaptations in POMC neurons, shifting the release from *α*-MSH to β-EP (Koch et al., 2015). In view of our current findings, it is possible that mTORC1 regulates POMC processing in response to different endogenous or exogenous stimuli, and that such processing might differ depending on the specific neurotransmitter cell subtype engaged in different moments of the feeding process.

## 5. Conclusions

The present study provides compelling evidence for a more sophisticated role of POMC neurons in the regulation of feeding than what we have so far believed, as specific subpopulations of these neurons may differentially impact this behavior. These findings reveal an unforeseen neuronal mechanism, which modifies our notion of hypothalamic circuits regulating food intake, energy balance and possibly their associated pathological states.

## Author Contributions

N.S., W.M., V.S., E.B., C.C., L.B., R.A.R., I.M., P.Z., S.L., C.Q., A.C., K.M. D.G., S.C. and

J.M.B. generated the data; N.S., W.M., V.S., G.S.H.Y., F.T.M., S.L.W., T.H., F.M. and D.C. analyzed the data together with the other authors; T.H., F.M. and G.M. critically contributed to discussion; D.C. conceptualized all studies and supervised the work; N.S., W.M., V.S., and

D.C. wrote the manuscript; all authors edited and approved the final version of the manuscript.

## Data statement

All data and resources related to the current study are available from the corresponding author upon reasonable request.

## Acknowledgements

We thank the animal, genotyping and bioinformatics platforms of the INSERM U1215, funded by INSERM and Labex Brain, for animal care, mouse lines management and genotyping. We thank the analytical chemistry facility of the INSERM U1215, funded by INSERM, for endocannabinoids quantification, and UT Southwestern and IGBMC for POMC-CreER^T2^ mice. The microscopy was done in the Bordeaux Imaging Center, a service unit of the CNRS-INSERM and Bordeaux University, member of the national infrastructure France BioImaging supported by the Labex Brain. The help of F. Cordelières, C. Poujol, S. Marais and P. Mascalchi (University of Bordeaux) is acknowledged. We thank the biochemistry and biophysics platform of Bordeaux NeuroCampus, supported by the Labex Brain, and Dr. A. Zeisel and Dr. S. Linnarsson (Karolinska Institutet, Stockholm, Sweden) for help with single-cell RNA sequencing studies. We thank Drs. S.C. Woods (University of Cincinnati), F. Chaouloff, and P. Ciofi (INSERM U1215) for useful suggestions, and C. Padgett for artwork. Supported by INSERM (D.C., C.Q., G.M.), Aquitaine Region, ANR-13-BSV4-0006, ANR-17-CE14-0029 and Labex BRAIN ANR-10-LABX-43 (D.C., G.M.); ANR-2010-1414-01, ANR-10-EQX-008- 1 OPTOPATH and FFRD (D.C.); ANR-16-CE37-0010 (G.M.); INSERM/Aquitaine Region PhD fellowship and FRM PhD fellowship FDT20150532545 (N.S.); PhD fellowship from the French Ministry for Higher Education, Research and Innovation and FRM PhD fellowship FDT201805005371 (V.S.); PhD fellowship from the French Ministry for Higher Education, Research and Innovation (S.L.); French Societies of Endocrinology (SFE) and Nutrition (SFN) (C.Q.); Marie Curie IRG n°224757 (D.C.), HEALTH-F2-2008-223713 and HEALTH-603191 (G.M.), FP7-People2009-IEF-251494 (D.C., E.B.), ERC-2010-StG-260515 and ERC-2014-PoC-640923 (G.M.), FRM (C.C., F.M., L.B., G.M.); Postgraduate Scholarship Programme of the Free State of Saxony and Erasmus Traineeship Programme (J.M.B.); F.T.M. is a New York Stem Cell Foundation – Robertson Investigator supported by the Medical Research Council (MR/P501967/1) and the Wellcome Trust and Royal Society (211221/Z/18/Z); NIH DK080003 (S.L.W.); Swedish Research Council, Hjärnfonden, the Novo Nordisk Foundation, ERC SECRET-CELLS, intramural funds of the Medical University of Vienna (T.H.); EMBO long-term research fellowship ALTF 596-2014 and Marie Curie Actions EMBOCOFUND2012 GA-2012-600394 (R.A.R.).

**Appendix A. Supplementary information.**

## References

1. Alexander, G.M., Rogan, S.C., Abbas, A.I., Armbruster, B.N., Pei, Y., Allen, J.A., et al., 2009. Remote control of neuronal activity in transgenic mice expressing evolved G protein-coupled receptors. Neuron. 63:27–39.

2. Aponte, Y., Atasoy, D., Sternson, S.M., 2011. AGRP neurons are sufficient to orchestrate feeding behavior rapidly and without training. Nat. Neurosci. 14:351–355.

3. Atasoy, D., Aponte, Y., Su, H.H., Sternson, S.M., 2008. A FLEX switch targets Channelrhodopsin-2 to multiple cell types for imaging and long-range circuit mapping. J. Neurosci. 28:7025–7030.

4. Balthasar, N., Coppari, R., McMinn, J., Liu, S.M, Lee, C.E., Tang, V., et al., 2004. Leptin receptor signaling in POMC neurons is required for normal body weight homeostasis. Neuron. 42:983–991.

5. Balthasar, N., Dalgaard, L.T., Lee, C.E., Yu, J., Funahashi, H., Williams, T., et al., 2005. Divergence of melanocortin pathways in the control of food intake and energy expenditure. Cell. 123:493–505.

6. Bellocchio, L., Lafenetre, P., Cannich, A., Cota, D., Puente, N., Grandes, P., et al., 2010. Bimodal control of stimulated food intake by the endocannabinoid system. Nat. Neurosci. 13:281–283.

7. Bellocchio, L., Soria-Gomez, E., Quarta, C., Metna-Laurent, M., Cardinal, P., Binder, E., et al., 2013. Activation of the sympathetic nervous system mediates hypophagic and anxiety-like effects of CB(1) receptor blockade. Proc. Natl. Acad. Sci. U. S. A. 110:4786–4791.

8. Berglund, E.D., Liu, C., Sohn, J.W., Liu, T., Kim, M.H., Lee, C.E., et al., 2013. Serotonin 2C receptors in pro-opiomelanocortin neurons regulate energy and glucose homeostasis. J. Clin. Invest. 123:5061–5070.

9. Betley, J.N., Cao, Z.F., Ritola, K.D., Sternson, S.M., 2013. Parallel, redundant circuit organization for homeostatic control of feeding behavior. Cell. 155:1337–1350.

10. Bockaert, J., Marin, P., 2015. mTOR in Brain Physiology and Pathologies. Physiol. Rev. 95:1157–1187.

11. Brandt, C., Nolte, H., Henschke, S., Engstrom, Ruud, L., Awazawa, M., Morgan, D.A., et al., 2018. Food Perception Primes Hepatic ER Homeostasis via Melanocortin-Dependent Control of mTOR Activation. Cell. 175:1321–1335 e1320.

12. Brown, L.M., Clegg, D.J., Benoit, S.C., Woods, S.C., 2006. Intraventricular insulin and leptin reduce food intake and body weight in C57BL/6J mice. Physiol. Behav. 89:687–691.

13. Burke, L.K., Darwish, T., Cavanaugh, A.R., Virtue, S., Roth, E., Morro, J., et al., 2017. mTORC1 in AGRP neurons integrates exteroceptive and interoceptive food-related cues in the modulation of adaptive energy expenditure in mice. Elife. 6.

14. Busquets-Garcia, A., Bains, J., Marsicano, G., 2018. CB_1_ Receptor Signaling in the Brain: Extracting Specificity from Ubiquity. Neuropsychopharmacology. 43:4–20.

15. Campbell, J.N., Macosko, E.Z., Fenselau, H., Pers, T.H., Lyubetskaya, A., Tenen, D., et al., 2017. A molecular census of arcuate hypothalamus and median eminence cell types. Nat. Neurosci. 20:484–496.

16. Caron, A., Labbe, S.M., Mouchiroud, M., Huard, R., Richard, D., Laplante, M., 2016. DEPTOR in POMC neurons affects liver metabolism but is dispensable for the regulation of energy balance. Am. J. Physiol. Regul. Integr. Comp. Physiol. 310:R1322–1331.

17. Castillo, P.E., Younts, T.J., Chavez, A.E., Hashimotodani, Y., 2012. Endocannabinoid signaling and synaptic function. Neuron. 76:70–81.

18. Chen, Y., Lin, Y.C., Kuo, T.W., Knight, Z.A., 2015. Sensory detection of food rapidly modulates arcuate feeding circuits. Cell. 160:829–841.

19. Cota, D., Proulx, K., Seeley, R.J., 2007. The role of CNS fuel sensing in energy and glucose regulation. Gastroenterology. 132:2158–2168.

20. Cota, D., Proulx, K., Smith, K.A., Kozma, S.C., Thomas, G., Woods, S.C., et al., 2006. Hypothalamic mTOR signaling regulates food intake. Science. 312:927–930.

21. Courtin, J., Chaudun, F., Rozeske, R.R., Karalis, N., Gonzalez-Campo, C., Wurtz, H., et al., 2014. Prefrontal parvalbumin interneurons shape neuronal activity to drive fear expression. Nature. 505:92–96.

22. Dagon, Y., Hur, E., Zheng, B., Wellenstein, K., Cantley, L.C., Kahn, B.B., 2012. p70S6 kinase phosphorylates AMPK on serine 491 to mediate leptin’s effect on food intake. Cell. Metab. 16:104–112.

23. Dennison, C.S., King, C.M., Dicken, M.S., Hentges, S.T., 2016. Age-dependent changes in amino acid phenotype and the role of glutamate release from hypothalamic proopiomelanocortin neurons. J. Comp. Neurol. 524:1222–1235.

24. Dicken, M.S., Tooker, R.E., Hentges, S.T., 2012. Regulation of GABA and glutamate release from proopiomelanocortin neuron terminals in intact hypothalamic networks. J. Neurosci. 32:4042–4048.

25. Dietrich, M.O., Horvath, T.L., 2013. Hypothalamic control of energy balance: insights into the role of synaptic plasticity. Trends Neurosci. 36:65–73.

26. Fan, W., Boston, B.A., Kesterson, R.A., Hruby, V.J., Cone, R.D., 1997. Role of melanocortinergic neurons in feeding and the agouti obesity syndrome. Nature. 385:165–168.

27. Fekete, C., Zseli, G., Singru, P.S., Kadar, A., Wittmann, G., Fuzesi, T., et al., 2012. Activation of Anorexigenic Pro-Opiomelanocortin Neurones during Refeeding is Independent of Vagal and Brainstem Inputs. J. Neuroendocrinol. 24:1423–1431.

28. Fenselau, H., Campbell, J.N., Verstegen, A.M., Madara, J.C., Xu, J., Shah, B.P., et al., 2017. A rapidly acting glutamatergic ARC-->PVH satiety circuit postsynaptically regulated by alpha-MSH. Nat. Neurosci. 20:42–51.

29. Gatta-Cherifi, B., Matias, I., Vallee, M., Tabarin, A., Marsicano, G., Piazza, P.V., et al., 2012. Simultaneous postprandial deregulation of the orexigenic endocannabinoid anandamide and the anorexigenic peptide YY in obesity. Int. J. Obes. (Lond) 36:880–885.

30. Haissaguerre, M., Saucisse, N., Cota, D., 2014. Influence of mTOR in energy and metabolic homeostasis. Mol. Cell. Endocrinol. 397:67–77.

31. Haissaguerre, M., Ferriere, A., Simon, V., Saucisse, N., Dupuy, N., Andre, C., et al., 2018. mTORC1-dependent increase in oxidative metabolism in POMC neurons regulates food intake and action of leptin. Mol. Metab. 12:98–106.

32. Hara, K., Maruki, Y., Long, X., Yoshino, K., Oshiro, N., Hidayat, S., et al., 2002. Raptor, a binding partner of target of rapamycin (TOR), mediates TOR action. Cell. 110:177–189.

33. Henry, F.E., Sugino, K., Tozer, A., Branco, T., Sternson, S.M., 2015. Cell type-specific transcriptomics of hypothalamic energy-sensing neuron responses to weight-loss. Elife 4.

34. Hentges, S.T., Otero-Corchon, V., Pennock, R.L., King, C.M., Low, M.J., 2009. Proopiomelanocortin expression in both GABA and glutamate neurons. J. Neurosci. 29:13684–13690.

35. Hentges, S.T., Nishiyama, M., Overstreet, L.S., Stenzel-Poore, M., Williams, J.T., Low, M.J., 2004. GABA release from proopiomelanocortin neurons. J. Neurosci. 24:1578–1583.

36. Huang da, W., Sherman, B.T., Lempicki, R.A., 2009. Systematic and integrative analysis of large gene lists using DAVID bioinformatics resources. Nat. Protoc. 4:44–57.

37. Jones, G.L., Wittmann, G., Yokosawa, E.B., Yu, H., Mercer, A.J., Lechan, R.M., et al., 2019. Selective Restoration of Pomc Expression in Glutamatergic POMC Neurons: Evidence for a Dynamic Hypothalamic Neurotransmitter Network. eNeuro. 6.

38. Kirkham, T.C., Williams, C.M., Fezza, F., Di Marzo, V., 2002. Endocannabinoid levels in rat limbic forebrain and hypothalamus in relation to fasting, feeding and satiation: stimulation of eating by 2-arachidonoyl glycerol. Br. J. Pharmacol. 136:550–557.

39. Koch, M., Varela, L., Kim, J.G., Kim, J.D., Hernandez-Nuno, F., Simonds, S.E., et al., 2015. Hypothalamic POMC neurons promote cannabinoid-induced feeding. Nature. 519:45–50.

40. Krashes, M.J., Lowell, B.B., Garfield, A.S., 2016. Melanocortin-4 receptor-regulated energy homeostasis. Nat. Neurosci. 19:206–219.

41. Lam, B.Y.H., Cimino, I., Polex-Wolf, J., Nicole Kohnke, S., Rimmington, D., Iyemere, V., et al., 2017. Heterogeneity of hypothalamic pro-opiomelanocortin-expressing neurons revealed by single-cell RNA sequencing. Mol. Metab. 6:383–392.

42. Luther, J.A., Tasker, J.G., 2000. Voltage-gated currents distinguish parvocellular from magnocellular neurones in the rat hypothalamic paraventricular nucleus. J. Physiol. 523 Pt 1:193–209.

43. Marsicano, G., Wotjak, C.T., Azad, S.C., Bisogno, T., Rammes, G., Cascio, M.G., et al., 2002. The endogenous cannabinoid system controls extinction of aversive memories. Nature. 418:530–534.

44. Marsicano, G., Goodenough, S., Monory, K., Hermann, H., Eder, M., Cannich, A., et al., 2003. CB_1_ cannabinoid receptors and on-demand defense against excitotoxicity. Science. 302:84–88.

45. Mazier, W., Saucisse, N., Gatta-Cherifi, B., Cota, D., 2015. The Endocannabinoid System: Pivotal Orchestrator of Obesity and Metabolic Disease. Trends Endocrinol. Metab. 26:524–537.

46. Mazier, W., Saucisse, N., Simon, V., Cannich, A., Marsicano, G., Massa F., et al., 2019. mTORC1 and CB_1_ receptor signaling regulate excitatory glutamatergic inputs onto the hypothalamic paraventricular nucleus in response to energy availability. Mol. Metab. 28:151–159.

47. Monge-Roffarello, B., Labbe, S.M., Roy, M.C., Lemay, M.L., Coneggo, E., Samson, P., et al., 2014. The PVH as a site of CB_1_-mediated stimulation of thermogenesis by MC4R agonism in male rats. Endocrinology. 155:3448–3458.

48. Monory, K., Massa, F., Egertová, M., Eder, M., Blaudzun, H., Westenbroek, R., et al., 2006. The endocannabinoid system controls key epileptogenic circuits in the hippocampus. Neuron. 51:455–466.

49. Mori, H., Inoki, K., Munzberg, H., Opland, D., Faouzi, M., Villanueva, E.C., et al., 2009. Critical role for hypothalamic mTOR activity in energy balance. Cell. Metab. 9:362–374.

50. Padilla, S.L., Carmody, J.S., Zeltser, L.M., 2010. Pomc-expressing progenitors give rise to antagonistic neuronal populations in hypothalamic feeding circuits. Nat. Med. 16:403–405.

51. Papadopoulos, A.D., Wardlaw, S.L., 1999. Endogenous alpha-MSH modulates the hypothalamic-pituitary-adrenal response to the cytokine interleukin-1beta. J. Neuroendocrinol. 11:315–319.

52. Reich, M., Liefeld, T., Gould, J., Lerner, J., Tamayo, P., Mesirov, J.P., 2006. GenePattern 2.0. Nat. Genet. 38:500–501.

53. Romanov, R.A., Zeisel, A., Bakker, J., Girach, F., Hellysaz, A., Tomer, R., et al., 2017. Molecular interrogation of hypothalamic organization reveals distinct dopamine neuronal subtypes. Nat. Neurosci. 20:176–188.

54. Sarbassov, D.D., Ali, S.M., Kim, D.H., Guertin, D.A., Latek, R.R., Erdjument-Bromage, H., et al., 2004. Rictor, a novel binding partner of mTOR, defines a rapamycin-insensitive and raptor-independent pathway that regulates the cytoskeleton. Curr. Biol. 14:1296–1302.

55. Savontaus, E., Breen, T.L., Kim, A., Yang, L.M., Chua, S.C., Jr., Wardlaw, S.L., 2004. Metabolic effects of transgenic melanocyte-stimulating hormone overexpression in lean and obese mice. Endocrinology. 145:3881–3891.

56. Saxton, R.A., Sabatini, D.M., 2017. mTOR Signaling in Growth, Metabolism, and Disease. Cell. 168:960–976.

57. Singru, P.S., Sanchez, E., Fekete, C., Lechan, R.M., 2007. Importance of melanocortin signaling in refeeding-induced neuronal activation and satiety. Endocrinology. 148:638–646.

58. Singru, P.S., Wittmann, G., Farkas, E., Zseli, G., Fekete, C., Lechan, R.M., 2012. Refeeding-activated glutamatergic neurons in the hypothalamic paraventricular nucleus (PVN) mediate effects of melanocortin signaling in the nucleus tractus solitarius (NTS). Endocrinology. 153:3804–3814.

59. Smith, M.A., Katsouri, L., Irvine, E.E., Hankir, M.K., Pedroni, S.M., Voshol, P.J., et al., 2015. Ribosomal S6K1 in POMC and AgRP Neurons Regulates Glucose Homeostasis but Not Feeding Behavior in Mice. Cell. Rep. 11:335–343.

60. Soria-Gomez, E., Massa, F., Bellocchio, L., Rueda-Orozco, P.E., Ciofi, P., Cota, D., et al., 2014a. Cannabinoid type-1 receptors in the paraventricular nucleus of the hypothalamus inhibit stimulated food intake. Neuroscience. 263:46–53.

61. Soria-Gomez, E., Bellocchio, L., Reguero, L., Lepousez, G., Martin, C., Bendahmane, M., et al., 2014b. The endocannabinoid system controls food intake via olfactory processes. Nat. Neurosci. 17:407–415.

62. Tong, Q., Ye, C.P., Jones, J.E., Elmquist, J.K., Lowell, B.B., 2008. Synaptic release of GABA by AgRP neurons is required for normal regulation of energy balance. Nat. Neurosci. 11:998–1000.

63. Tsigos, C., Crosby, S.R., Gibson, S., Young, R.J., White, A., 1993. Proopiomelanocortin is the predominant adrenocorticotropin-related peptide in human cerebrospinal fluid. J. Clin. Endocrinol. Metab. 76:620–624.

64. Verty, A.N., McFarlane, J.R., McGregor, I.S., Mallet, P.E., 2004. Evidence for an interaction between CB_1_ cannabinoid and melanocortin MCR-4 receptors in regulating food intake. Endocrinology. 145:3224–3231.

65. Wardlaw, S.L., 2011. Hypothalamic proopiomelanocortin processing and the regulation of energy balance. Eur. J. Pharmacol. 660:213–219.

66. Weston, M.C., Chen, H., Swann, J.W., 2012. Multiple roles for mammalian target of rapamycin signaling in both glutamatergic and GABAergic synaptic transmission. J. Neurosci. 32:11441–11452.

67. Wittmann, G., Hrabovszky, E., Lechan, R.M., 2013. Distinct glutamatergic and GABAergic subsets of hypothalamic pro-opiomelanocortin neurons revealed by in situ hybridization in male rats and mice. J. Comp. Neurol. 521:3287–3302.

68. Yang, S.B., Tien, A.C., Boddupalli, G., Xu, A.W., Jan, Y.N., Jan, L.Y., 2012. Rapamycin ameliorates age-dependent obesity associated with increased mTOR signaling in hypothalamic POMC neurons. Neuron. 75:425–436.

69. Zhan, C., Zhou, J., Feng, Q., Zhang, J.E., Lin, S., Bao, J., et al., 2013. Acute and long-term suppression of feeding behavior by POMC neurons in the brainstem and hypothalamus, respectively. J. Neurosci. 33:3624–3632.

