## Supplemental Material and Tables for "POMC neurons functional heterogeneity relies on mTORC1 signaling"

***Detailed protocol of the Triple-FISH combined with IHC.*** Brain sections from C57BL/6J mice were processed for the co-expression of POMC, vGlut2, GAD65, GAD67 and c-Fos, to evaluate the phenotype of activated POMC neurons after short-term exposure to a palatable diet. We took 4 free-floating sections per animal containing POMC neurons and spaced 300µm apart, in order to have an appropriate rostro-caudal representation of the ARC. Day 1: After 3 washes in PBS, endogenous peroxidases were quenched in 3% H2O2 in PBST (PBS+0.1% Tween20) for 30min. After 1 wash in PBS, endogenous biotins were blocked by incubating the slices in Avidin solution for 15min. They were then washed in PBS, incubated in Biotin solution for another 15min (Avdin/Biotin blocking kit, Vector) and washed again in PBS. Rabbit anti-cFos antibody (Cell Signaling #2250, 1/1000) in PBS+0.3% Triton X-100 was incubated overnight at 4°C. Day 2: Sections were washed 3 times in PBS, incubated in HRP-conjugated goat anti-rabbit (Cell Signaling #7074, 1/500) in PBS+0.3% Triton X-100 for 2h and washed 3 times again with PBS. Tyramide signal amplification (TSA, AkoyaBio) was done, using tyramine-labeled biotins (TSA-Biotin dissolved in 1X Plus Amplification buffer, 1/250, 30min). After 3 washes in PBS, HRP were blocked by a 30min-incubation in 3% H2O2 in PBS followed by a 20min-incubation in 0.2M HCl (with 1 wash in PBS in between). Biotins were fixed by a 10min-incubation in ice-cold 10% NBF. After 1 wash in PBS-DEPC (PBS with 0.01% diethylpyrocarbonate) and 2 washes in PBST-DEPC (PBS-DEPC+0.1% Tween20), sections were acetylated with a treatment of 0.25% acetic anhydride in 0.1M triethanolamine (pH=8.0) for 2x5min. After 1 wash in PBST-DEPC, the hybridization solution was prepared as follows: POMC-DNP, vGlut2-DIG, GAD65-FITC and GAD67-FITC probes (1/1000 for each of them) were dissolved in hybmix (50% deionized formamide, 20mM Tris at pH=8.0, 300mM NaCl, 5mM EDTA, 10% dextran sulfate, 1X Denhardt's solution, 0.5mg/mL tRNA, 0.2mg/mL acid-cleaved carrier DNA from salmon's sperm, 1M DTT dissolved in water containing 0.01% DEPC) and heated at 90°C for 5min to ensure probe linearization. Sections were then incubated in this hybmix overnight (16h-20h) at 70°C in a water bath. Day 3: A series of washes with increased stringency was carried out as follows:5X SSC (Saline Sodium Citrate) + 0.2% Tween20 for 5min at 65°C; 2X SSC + 0.2% Tween20 + 50% Formamide Amide for 30min at 65°C; 1X SSC + 0.2% Tween20 + 50% Formamide Amide for 30min at 65°C; 0.1X SSC + 0.2% Tween20 for 30min at 65°C. After the last stringent wash, sections were washed once with TNT buffer (100mM Tris at pH=7.5 + 150mM NaCl + 0.05% Tween20), incubated 30min with 3% H2O2 in PBS followed by a 20min-incubation in 0.2M HCl (with 1 wash in TNT in between). After a wash in TNT, sections were incubated for 1h in blocking TNB buffer (AkoyaBio, prepared as per manufacturer's instructions) then overnight at 4°C in HRP-labeled anti-DIG antibody (Sigma #11207733910, 1/1500) in TNB. Day 4: Sections were washed 3 times in TNT, and subjected to TSA-Cy3 revelation (TSA-Cy3 dissolved in 1X Plus Amplification buffer, 1/100, 30min; AkoyaBio). After 3 washes in TNT, HRP were blocked by a 30min-incubation in 3% H2O2 in PBS followed by a 20min-incubation in 0.2M HCl (with 1 wash in TNT in between). After 1 wash in TNT, sections were incubated in HRP-conjugated anti-FITC antibody (Sigma #11426346910, 1/1500) in TNB for 2h. Sections were washed 3 times in TNT and subjected to TSA-FITC revelation (TSA-FITC dissolved in 1X Plus Amplification buffer, 1/100, 30min, AkoyaBio). After 3 washes in TNT, HRP were blocked by a 30min-incubation in 3% H2O2 in PBS followed by a 20min-incubation in 0.2M HCl (with 1 wash in TNT in between). After 1 wash in TNT, sections were incubated in HRP-conjugated anti-DNP antibody (AkoyaBio, FP1129, 1/100) in TNB overnight at 4°C. Day 5: Sections were washed 3 times in TNT, and subjected to bench-made TSA-Cy5 revelation (bench-made TSA-Cy5 reagent dissolved in bench-made Amplification buffer, 1/200, 30min). After 3 washes in TNT, HRP were blocked by a 30min-incubation in 3% $H_2O_2$ in PBS followed by a 20min-incubation in 0.2M HCl (with 1 wash in TNT in between). After 1 wash in TNT, sections were incubated for 30min in Avidin-Biotin complexes (ABC, Vector) prepared as per manufacturer's instruction (1/50 of solution A and 1/50 of solution B in PBST, mixed at least 30min before the incubation). Sections were then washed 3 times in TNT and subjected to TSA-Coumarin revelation (TSA-Coumarin dissolved in 1X Plus Amplification buffer, 1/100, 30min, AkoyaBio). After 3 washes in TNT, HRP were blocked by a 30min-incubation in 3% $H_2O_2$ in PBS followed by a 20min-incubation in 0.2M HCl (with 1 wash in TNT in between). After 2 washes in TNT and 1 final wash in 50mM Tris (pH=7.5), sections were mounted using ProLong Gold medium, cover-slipped (0.13-0.17mm thickness) and stored in the dark.

**Bench-made TSA-Cy5 reagents.** First, Cy5-NHS ester (Amersham) was dissolved in N,N-dimethylformamide (DMF) to obtain a 10mg/mL solution. In parallel, a 10mg/mL tyramine solution was prepared by dissolving tyramine in [DMF+1% triethylamine]. These two solutions were then mixed at a 1:1 molar ratio and the mixture was incubated in the dark for 2h. Finally, an appropriate amount of 100% ethanol was added to get a final concentration of 5mg/mL of Cy5 after which aliquots were made and stored at -20°C. Bench-made amplification buffer was produced as follows: a stock solution was prepared by dissolving 2% dextran sulfate, 0.1% Tween20 and 500µg/mL of 4-iodophenol in 0.1M borate solution and stored at 4°C. Then, 0.006% of $H_2O_2$ was added extemporaneously to make the amplification buffer.

**Table S1.** List of primers used for mouse genotyping by PCR.

| Mouse line | Primer | sequence 5'3' | expected fragment & length | |
|---|---|---|---|---|
| POMC-Cre | Cre-F | GCGGTCTGGCAGTAAAAACTATC | Cre | 100 pb |
| | Cre-R | GTGAAACAGCATTGCTGTCACTT | | |
| POMC-CreER$^{T2}$ | Cre-F | GCGGTCTGGCAGTAAAAACTATC | CreER$^{T2}$ | 100 bp |
| | Cre-R | GTGAAACAGCATTGCTGTCACTT | | |
| $CB_1$-Flox | $CB_1$-F | GCTGTCTCTGGTCCTCTTAAA | wt | 450 bp |
| | $CB_1$-R | GGTGTCACCTCTGAAAACAGA | Flox | 555 bp |
| | $CB_1$-ex | CTCCTGTATGCCATAGCTCTT | excision | 680 bp |
| Rictor-Flox | Rictor-F | CAAGCATCATGCAGCTCTTC | Flox | 560 bp |
| | Rictor-R | TCCCAGAATTTCCAGGCTTA | wt | 410 bp |
| Rptor-Flox | Rptor-F | CTCAGTAGTGGTATGTGCTCAG | Flox | 200 bp |
| | Rptor-R | GGGTACAGTATGTCAGCACAG | wt | 160 bp |
| vGlut2-flox | vglut2-F | CTGAGCGAAGGTGAGCTGAA | Flox | 400 bp |
| | vglut2-R | TGGGCCAGAACACAGGATATG | wt | 270 bp |
| | vglut2-ex | TCACTGCCTTGTTCCTAGTGC | excision | 700 bp |
| POMC-YFP | WT-F | AAAGTCGCTCTGAGTTGTTAT | wt | 620 bp |
| | Ki-F | GCGAAGAGTTTGTCCTCAACC | Ki | 330 bp |
| | Common-R | GGAGCGGGAGAAATGGATATG | | |
| Ai6 | Ai6-WT-F | AAGGGAGCTGCAGTGGAGTA | wt | 300 bp |
| | Ai6-WT-R | CCGAAAATCTGTGGGAAGTC | | |
| | Ai6-mut-F | AACCAGAAGTGGCACCTGAC | Ai6 | 210 bp |
| | Ai6-mut-R | GGCATTAAAGCAGCGTATCC | | |

**Table S2.** Detailed data about the different light-evoked POMC inputs recorded in 17 connected parvocellular neurons (related to Figure 3). Recorded cell features (Em, Cm and Rm) were collected right after patch establishment. Parameters on light-evoked transmission (Peak amplitude, Time of peak and Decay time constant) are indicated as the mean of 4 minutes (± sem) after at least 10 minutes of treatment.

| Recorded cell | Treatment | Peak Amplitude (pA) | Time of peak (msec) | Decay (msec) |
|---|---|---|---|---|
| ***Pure Glut (no effect of picrotoxin)*** | | | | |
| Cell# 1 | Veh | -32.3 ± 3.41 | 15.1 ± 0.16 | 2.7 ± 0.23 |
| -44mV 22pF MΩ | Picrotoxin | -33.1 ± 0.47 | 14.9 ± 0.05 | 2.6 ± 0.18 |
| Cell# 2 | Veh | -43.8 ± 2.62 | 19.8 ± 0.1 | 2.3 ± 0.08 |
| -43mV 19pF 750MΩ | Picrotoxin | -46.3 ± 1.43 | 21.5 ± 0.53 | 2.0 ± 0.11 |
| Cell# 3 | Veh | -24.5 ± 1.32 | 14.8 ± 0.08 | 2.6 ± 0.17 |
| -44mV 22pF 750MΩ | Picrotoxin | -27.9 ± 2.02 | 15.1 ± 0.15 | 2.4 ± 0.13 |
| Cell# 4 | Veh | -138.0 ± 0.97 | 12.5 ± 0.03 | 3.5 ± 0.04 |
| -52mV 34pF 750MΩ | Picrotoxin | -144.0 ± 1.70 | 12.5 ± 0.02 | 3.2 ± 0.09 |
| ***Pure Glut (full blockade by NBQX APV)*** | | | | |
| Cell# 5 | Veh | -22.8 ± 1.09 | 16.1 ± 0.30 | 2.7 ± 0.20 |
| -42mV 23pF 800MΩ | NBQX/APV | no peak | no peak | no peak |
| Cell# 6 | Veh | -32.4 ± 1.06 | 10.9 ± 0.04 | 1.6 ± 0.01 |
| -48mV 23pF 710MΩ | NBQX/APV | no peak | no peak | no peak |
| Cell# 7 | Veh | -27.5 ± 0.91 | 13.1 ± 0.07 | 1.9 ± 0.34 |
| -49mV 26pF 650MΩ | NBQX/APV | no peak | no peak | no peak |
| Cell# 8 | Veh | -26.0 ± 1.52 | 11.6 ± 0.03 | 2.2 ± 0.04 |
| -46mV 21pF 850MΩ | NBQX/APV | no peak | no peak | no peak |
| Cell# 9 | Veh | -87.3 ± 2.44 | 21.5 ± 0.18 | 3.1 ± 0.05 |
| -49mV 18pF 750MΩ | NBQX/APV | no peak | no peak | no peak |
| Cell# 10 | Veh | -40.2 ± 2.21 | 12.5 ± 0.23 | 2.1 ± 0.17 |
| -46mV 23pF 930MΩ | NBQX/APV | no peak | no peak | no peak |
| ***Pure GABA (full blockade by picrotoxin)*** | | | | |
| Cell# 11 | Veh | -79.0 ± 1.44 | 50.6 ± 0.63 | 7.6 ± 0.16 |
| -44mV 19pF 1500MΩ | Picrotoxin | no peak | no peak | no peak |
| Cell# 12 | Veh | -56.0 ± 1.78 | 15.1 ± 0.09 | 11.7 ± 0.67 |
| -42mV 15pF 2000MΩ | Picrotoxin | no peak | no peak | no peak |
| Cell# 13 | Veh | -25.7 ± 0.82 | 18.7 ± 0.29 | 8.1 ± 0.64 |
| -44mV 17pF 2000MΩ | Picrotoxin | no peak | no peak | no peak |
| ***GABA/Glut (isolation of Glut)*** | | | | |
| Cell# 14 | Veh | -220.9 ± 7.31 | 14.3 ± 0.04 | 6.3 ± 0.21 |
| -42mV 17pF 900MΩ | Picrotoxin | -26.7 ± 1.09 | 12.4 ± 0.06 | 2.9 ± 0.27 |
| Cell# 15 | Veh | -20.7 ± 0.25 | 18.3 ± 0.07 | 3.5 ± 0.07 |
| -48mV 22pF 820MΩ | Picrotoxin | -24.9 ± 1.21 | 18.3 ± 0.52 | 2.9 ± 0.14 |
| ***GABA/Glut (isolation of GABA)*** | | | | |
| Cell# 16 | Veh | -72.7 ± 2.14 | 13.3 ± 0.03 | 5.4 ± 0.07 |
| -45mV 19pF 750MΩ | NBQX/APV | -44.9 ± 0.96 | 13.3 ± 0.06 | 5.9 ± 0.17 |
| Cell# 17 | Veh | -63.7 ± 1.20 | 20.7 ± 0.08 | 5.1 ± 0.12 |
| -48mV 23pF 1800MΩ | NBQX/APV | -47.4 ± 2.24 | 21.7 ± 0.16 | 4.9 ± 0.04 |

**Table S3.** POMC and POMC-derived peptides ACTH and β-EP levels in the hypothalamus and endocannabinoid levels in the cortex and the hippocampus of mice fasted or refed and treated with an icv injection of RAPA or its vehicle (related to Figure 5). Data are Mean ± SEM.

| Analyte | Fasted/Veh | Refed/Veh | Refed/RAPA | N mice | Analysis | Degree of freedom & F/t/R etc value | p-value |
|---|---|---|---|---|---|---|---|
| POMC (fmol/mg) | 254.18 ± 14.85 | 292.94 ± 29.63 | 281.99 ± 24.07 | 8 | One-way ANOVA | F (2, 21) = 0.7140 | 0.5012 |
| ACTH (fmol/mg) | 71.98 ± 11.06 | 84.88 ± 12.95 | 78.91 ± 8.15 | 8 | One-way ANOVA | F (2, 21) = 0.3511 | 0.7080 |
| β-EP (fmol/mg) | 462.2 ± 26.11 | 620.0 ± 88.03 | 502.4 ± 22.15 | 8 | One-way ANOVA | F (2, 21) = 2.261 | 0.1291 |
| AEA (fmol/mg) | | | | | | | |
| *Hippocampus* | 38.99 ± 4.99 | 43.14 ± 4.45 | 38.16 ± 4.03 | 3-5 | One-way ANOVA | F (2, 10) = 0.3972 | 0.6823 |
| *Cortex* | 35.22 ± 4.52 | 38.72 ± 4.38 | 38.25 ± 6.72 | 3-5 | One-way ANOVA | F (2, 10) = 0.08910 | 0.9155 |
| 2-AG (pmol/mg) | | | | | | | |
| *Hippocampus* | 12.55 ± 2.94 | 14.89 ± 1.14 | 13.85 ± 1.66 | 3-5 | One-way ANOVA | F (2, 10) = 0.3893 | 0.6873 |
| *Cortex* | 16.74 ± 0.69 | 15.38 ± 0.68 | 16.53 ± 2.15 | 3-5 | One-way ANOVA | F (2, 10) = 0.2237 | 0.8035 |

**Table S4**. Statistical analyses for main figures.

| Figure | Dependent variable(s) | Factor Analyzed | Analysis | n | Degree of freedom & F/t/R etc value | P value |
|---|---|---|---|---|---|---|
| 1B | Distance from ventricle | Group | Kruskal-Wallis ANOVA | 3914 cells | | p<0.0001 |
| 1C | Slope of the lines | Group | Linear regression | 3914 cells | F=1.185e+014 dfn=3 dfd=592 | p<0.0001 |
| 2B | POMC AP firing | Treatment | Paired t-test | 41 cells | t=7.175 df=40 | p<0.0001 |
| 2C | POMC AP firing | Treatment | Paired t-test | 37 cells | t=6.707 df=36 | p<0.0001 |
| 2D | POMC AP firing | Treatment | Paired t-test | 39 cells | t=1.692 df=38 | p=0.0989 |
| 2E | POMC cell Em | Group | Unpaired t-test | 41-37 cells | t=3.425 df=76 | p=0.0010 |
| 2F | POMC cell Rm | Group | Unpaired t-test | 41-37 cells | t=6.653 df=76 | p<0.0001 |
| 2G | POMC cell Cm | Group | Unpaired t-test | 41-37 cells | t=5.282 df=76 | p<0.0001 |
| 2H | RAPA effect vs POMC cell Cm | Correlation | Linear Regression | 78 cells | $R^2$=0.1191 | p=0.0020 |
| 2J | Ratio GAD spots number / Cm | Group | Unpaired t-test | 7-11 cells | t=3.063 df=16 | p=0.0074 |
| 2K | AP firing change (%) vs GAD spots number | Correlation | Linear Regression | 18 cells | $R^2$=0.6683 | p<0.0001 |
| 3D | % Change in eEPSC amplitude | Treatment | Paired t-test, one-tailed | 6 cells | t=6.802, df=5 | p=0.0005 |
| 3E | % Change in eIPSC amplitude | Treatment | Paired t-test, one-tailed | 5 cells | t=2.290, df=4 | p=0.041 |
| 4A | Food intake | Treatment | RM Two-way ANOVA | 5 mice | F (1, 8) = 2.295 | p=0.1682 |
| 4A | Food intake | Time | RM Two-way ANOVA | 5 mice | F (1,614, 12,91) = 53.85 | p<0.0001 |
| 4A | Food intake | Interaction | RM Two-way ANOVA | 5 mice | F (2, 16) = 11.17 | p=0.0009 |
| 4B | % c-Fos positive POMC neurons | Treatment | One-way ANOVA | 4-5 | F (2, 11) = 17.84 | p=0.0004 |
| C | % of p-S6 in c-Fos positive POMC neurons | Treatment | One-way ANOVA | 4-5 | F (2, 11) = 11.13 | p=0.0023 |
| 4E | 2h food intake | Genotype | RM Two-way ANOVA | 6-9 mice | F (1, 13) = 1.828 | p=0.1994 |
| 4E | 2h food intake | Treatment | RM Two-way ANOVA | 6-9 mice | F (1, 13) = 21.71 | p=0.0004 |
| 4E | 2h food intake | Interaction | RM Two-way ANOVA | 6-9 mice | F (1, 13) = 5.870 | p=0.0307 |
| 4E | 2h food intake | Matched individuals | RM Two-way ANOVA | 6-9 mice | F (13, 13) = 1,623 | p=0.1970 |
| 4F | 2h food intake | Genotype | RM Two-way ANOVA | 5 mice | F (1, 8) = 4.325 | p=0.0711 |
| 4F | 2h food intake | Treatment | RM Two-way ANOVA | 5 mice | F (1, 8) = 6,337 | p=0.0360 |

| | | | | | | |
|---|---|---|---|---|---|---|
| 4F | 2h food intake | Interaction | RM Two-way ANOVA | 5 mice | F (1, 8) = 0.6067 | p=0.4585 |
| 4F | 2h food intake | Matched individuals | RM Two-way ANOVA | 5 mice | F (8, 8) = 0.5056 | p=0.8229 |
| 4G | 2h food intake | Treatment | RM one-way ANOVA | 5 mice | F (1.938, 7.751) = 30,42 | p=0.0002 |
| 4G | 2h food intake | Matched individuals | RM one-way ANOVA | 5 mice | F (4, 8) = 7.995 | p=0.0067 |
| 5B | α-MSH levels | Treatment | One-way ANOVA | 8 mice | F (2, 21) = 4.440 | p=0.0246 |
| 5C | α-MSH/POMC | Treatment | Unpaired t-test | 8 mice | t=2.873 df=14 | p=0.0123 |
| 5D | α-MSH/ACTH | Treatment | Unpaired t-test | 8 mice | t=2.646 df=14 | p=0.0192 |
| 5E | α-MSH/β-EP | Treatment | Unpaired t-test | 8 mice | t=2.807 df=14 | p=0.0140 |
| 5F | AEA levels | Treatment | One-way ANOVA | 3-5 mice | F (2, 10) = 4.737 | p=0.0357 |
| 5G | 2-AG levels | Treatment | One-way ANOVA | 3-5 mice | F (2, 10) = 0,2019 | p=0.8204 |
| 5H | AEA levels | Treatment | Unpaired t-test | 6 mice | t=2.556 df=10 | p=0.0286 |
| 5I | AEA levels | Treatment | Unpaired t-test | 8-10 mice | t=2.171 df=16 | p=0.0453 |
| 6A | 2h food intake | Genotype | Two-way ANOVA | 8-18 mice | F (1, 78) = 0.111 | p=0.739 |
| 6A | 2h food intake | Treatment | Two-way ANOVA | 8-18 mice | F (2, 78) = 20.37 | p<0.0001 |
| 6A | 2h food intake | Interaction | Two-way ANOVA | 8-18 mice | F (2, 78) = 6.288 | p=0.0029 |
| 6B | 1h food intake | Genotype | Unpaired t-test | 8-14 mice | t=2.902 df=20 | p=0.0088 |
| 6C | 2h food intake | Treatment | Paired t-test | 8 mice | t=0.1839 df=7 | p=0.859 |

**Table S5**. Statistical analyses for supplemental figures.

| Figure | Dependent variable(s) | Factor Analyzed | Analysis | n | Degree of freedom & F/t/R etc value | P value |
|---|---|---|---|---|---|---|
| S3A | POMC AP firing | Treatment | Paired t-test | 117 cells | t=0.0793 df=116 | p=0.9369 |
| S3B | Change in Em under RAPA | Group | Unpaired t-test | 41-37 cells | t=5.244 df=76 | p<0.0001 |
| S3E | GAD staining vs POMC size | Correlation | Linear Regression | 18 cells | $R^2$=0.4008 | p=0.0048 |
| S3F | AP firing change (%) vs POMC size | Correlation | Linear Regression | 18 cells | $R^2$=0.5256 | p=0.0007 |
| S5B | % of Rictor expression in POMC neurons | Genotype | Unpaired t-test | 4 mice | t=61.38 df=6 | p<0.0001 |
| S5C | 24h food intake | Genotype | Unpaired t-test | 5 mice | t=1.124 df=8 | p=0.2937 |
| S5E | % c-Fos in mCherry-POMC positive neurons | Treatment | Unpaired t-test | 2-4 mice | t=3.774 df=4 | p=0.019 |
| S5F | % of p-S6 in mCherry and c-Fos positive POMC neurons | Treatment | Unpaired t-test | 2-4 mice | t=46.60 df=4 | p<0.0001 |
| S5G | 2h food intake | Treatment | Paired t-test | 6 mice | t=0.03704 df=5 | p=0.9719 |
| S5H | 2h food intake | Treatment | Paired t-test | 7 mice | t=3.090 df=6 | p=0.0214 |
| S6A | 2h food intake | Treatment | One-way ANOVA | 6-18 mice | F (7, 62) = 7.326 | p<0.0001 |
| S6B | AEA levels | Treatment | Unpaired t-test | 6 mice | t=0.183 df=10 | p=0.8584 |
| S7A | 24h food intake | Genotype | Unpaired t-test | 5-11 mice | t=0.1775 df=14 | p=0.8617 |
| S7B | 2h food intake | Treatment | One-way ANOVA | 6-8 mice | F (3, 25) = 0.770 | p=0.5217 |
| S7C | 24h food intake | Treatment | One-way ANOVA | 6-8 mice | F (3, 25) = 5.000 | p=0.0075 |
| S7D | mIPSC frequency | Treatment | Paired t-test | 10 cells | t=2.609 df=9 | p=0.0283 |
| S7E | mIPSC frequency | Treatment | Paired t-test | 12 cells | t=1.511 df=11 | p=0.159 |
| S7F | Change mIPSC | Genotype | Unpaired t-test | 10-12 cells | t=2.272 df=20 | p=0.0343 |
| S8A | 2h food intake | Diet | Mann-Whitney test | 4 mice | | p=0.0286 |
| S8B | c-Fos expression | Diet | Mann-Whitney test | 4 mice | | p=0.029 |
| S8D | c-Fos expression | Group | Kruskal-Wallis ANOVA | 4 mice | | p<0.0001 |
| S9C | 24h food intake | Genotype | Unpaired t-test | 15-8 mice | t=1.755 df=21 | p=0.0938 |

### Supplemental Figure Legends

**Figure S1. Analysis of the distribution of the different POMC subpopulations in the hypothalamic ARC.** (**A**) Rostro-caudal distribution of different POMC subpopulations in C57BL/6J mice (n=4), expressing or not GABAergic and glutamatergic markers. (**B**) Representative FISH images illustrating AgRP and POMC mRNA colocalization in the ARC of adult POMC-CreER$^{T2}$-Ai6 mice (n=2 mice). Scale bar in A is 100μm, scale bar in B is 50μm and 25μm for smaller inset. V: 3$^{rd}$ ventricle.

**Figure S2. POMC GABAergic, glutamatergic and GABA/glutamatergic neurons have specific gene expression profile.** Neurotransmitter type-specific single cell transcriptome profiling of hypothalamic *Pomc*$^+$ neurons. The heat map illustrates top rank 100 genes differently expressed in each *Pomc*$^+$ neurotransmitter-determined group (GABA, GLUT and GABA/GLUT, indicated by green rectangle). Left to heatmaps – p values for log transformed data, group of interest vs other 2 groups; right – gene names.

**Figure S3. Differential response of POMC neurons to mTORC1 blockade depending on their neurotransmitter type.** (**A**) Taking all analyzed POMC neurons together (n=117 cells), RAPA perfusion has no significant effect on cell firing (paired t-test). (**B**) Effect of RAPA treatment on cell membrane potential in POMC RAPA$^{act}$ neurons (+2.13 ± 0.43 mV, n=37 cells) and in POMC RAPA$^{inh}$ neurons (-1.23 ± 0.46 mV, n=41 cells); unpaired t-test. (**C**,**D**) Representative images (Z-stack 3D-reconstruction) of a recorded POMC neuron (C) and a hippocampal CA1 pyramidal cell (D). When running the immunofluorescence protocol in absence of the primary antibody anti-GAD65/67, POMC cells do not exhibit GAD signal within the cell surface (C), some background signal can still be observed; CA1 pyramidal cells (glutamatergic cells), which were used as negative control, do not have GAD staining (D). (**E**,**F**) POMC neurons capacitance negatively correlates with GAD spots quantity (E) and with the change in action potential firing induced by RAPA perfusion (F) (n= 18 cells, linear regression). Scale bar in C and D is 3μm. ***p<0.001. See also Table S5.

**Figure S4. Pharmacological strategies used to characterize the nature of light-evoked POMC inputs on parvocellular neurons.** Neurotransmission classification (mixed GABA/Glut, pure GABA and pure Glut) was made after perfusion of picrotoxin to block $GABA_A$ related currents (**A**) or of NBQX/APV to block AMPA/NMDA receptors mediated currents (**B**).

**Figure S5. Expression of rictor protein in hypothalamic POMC neurons and characterization of DREADD strategy to activate hypothalamic POMC neurons.** (**A, B**) Representative images illustrating rictor (red) protein staining in POMC-expressing neurons (green) in the ARC of POMC-*Rictor*-KO and control littermates (A) and related quantification (B) (n=4 mice per group). Scale bar in A 50 μm; 5 μm for smaller inset. (**C**) Daily food intake in POMC-*Rictor*-KO and control littermates (n=5 mice per group). (**D**) POMC neurons in the ARC (green) express mCherry-labelled DREADD (red, arrowheads). Scale bar in D: 25 μm. (**E-G**) Representative images (E) from the ARC of POMC-$ARC^{hM3Dq}$ mice treated with CNO or its vehicle and related quantification of c-Fos expression (green, F) and phosphorylation of S6 (p-S6, purple, G) in POMC mCherry-positive neurons (red) (n=2-4 mice per group). Scale bar in E: 50 μm; 10 μm for smaller inset. (**H**) Acute ip injection of CNO does not alter refeeding response in POMC-$ARC^{hM3Dq}$ mice having received an icv administration of DMSO (vehicle used for RAPA) (n=6 mice per group). (**I**) Acute ip CNO injection does not alter RAPA-induced hyperphagia in POMC-ARC$^{hM3Dq}$-control littermates that do not express the Cre recombinase (n=7 mice per group). Data analyzed by unpaired t-test (B, C, F, G) or paired t-test (H, I). *p<0.05 and ***p<0.001. V: 3$^{rd}$ ventricle. See also Table S5.

**Figure S6. Effect of the α-MSH analog MTII on food intake and hypothalamic AEA levels.** (**A**) Dose-response effect of an icv administration of MTII on 2h refed C57BL/6J mice (n=6-18 mice per group, one-way ANOVA followed by Fisher LSD post-hoc test, 3 experiments combined together). (**B**) Hypothalamic AEA content in refed C57BL/6J mice treated with MTII (0.02 μg, icv) or its vehicle (n=6 mice per group, unpaired t-test). *p<0.05; **p<0.01; ***p<0.001. See also Table S5.

**Figure S7. Basal food intake and GABAergic transmission in POMC-*CB₁*-KO mice.** (**A**) Daily food intake in POMC-$CB_1$-KO and Control littermates (n=5-11 mice per group, unpaired t-test) (**B**, **C**) Dose-response effect of an icv administration of the $GABA_A$ receptor antagonist picrotoxin (Ptx) in refed C57BL/6J mice (n=6-8 mice per group, one-way ANOVA followed by Fisher LSD post-hoc test). (**D**-**F**) Effect of the $CB_1R$ agonist WIN55-212 on miniatures inhibitory post-synaptic currents (mIPSC) frequency of parvocellular neurons of the PVN of POMC-$CB_1$-Control (D, n=10 cells) and -KO littermates (E, n=12 cells); (F) Change in mIPC frequency under WIN (n=10-12 cells). Data analyzed by paired t-test (D, E) and unpaired t-test (F). *p<0.05, **p<0.01. See also Table S5.

**Figure S8. Acute exposure to palatable food preferentially recruits POMC/GABA neurons.** (**A**) 2h food intake during the light phase in chow-fed and palatable food-fed C57BL/6J mice (n=4 mice per group, Mann-Whitney test). (**B**) c-Fos expression in POMC neurons of the ARC of C57BL/6J mice fed chow or palatable food for 2h during the light phase (n=4 mice per group, Mann-Whitney test). (**C**) Representative images of simultaneous detection of POMC (blue), GAD65/67 (green) and vGlut2 (red) mRNA by FISH combined with immunodetection of c-Fos (white) in the ARC of C57BL/6J mice exposed to palatable food, and related quantification (**D**) of each POMC neurons subpopulation among all POMC-cFos+ neurons (n=4 mice, Kruskal-Wallis ANOVA followed by Dunn's post-test). Scale bar in C: 100µm; V= 3$^{rd}$ ventricle. *$p<0.05$, **$p<0.01$. See also Table S5.

**Figure S9. Evaluation of genetic deletion of vGlut2 in POMC neurons and basal food intake in POMC-*vGlut2*-KO mice.** (**A**) Representative images of the co-expression of Zsgreen1 (green) and POMC (red) in POMC-CreER$^{T2}$-Ai6 mice to demonstrate appropriate recombination in the ARC after tamoxifen (TAM) administration (n=4 mice per group). (**B**) Representative PCR showing deletion of vGlut2 in hypothalami of POMC-*vGlut2*-KO mice. Note the excision fragment obtained at 700bp in the KO after tamoxifen administration. (**C**) Mean daily food intake in POMC-*vGlut2*-KO and Control littermates (n=8-15 mice per group, unpaired t-test). Scale bar in A: 100µm; V= 3$^{rd}$ ventricle. See also Table S5.
