## Supplementary figures and images for "POMC neurons functional heterogeneity relies on mTORC1 signaling"

### Supplemental Figures

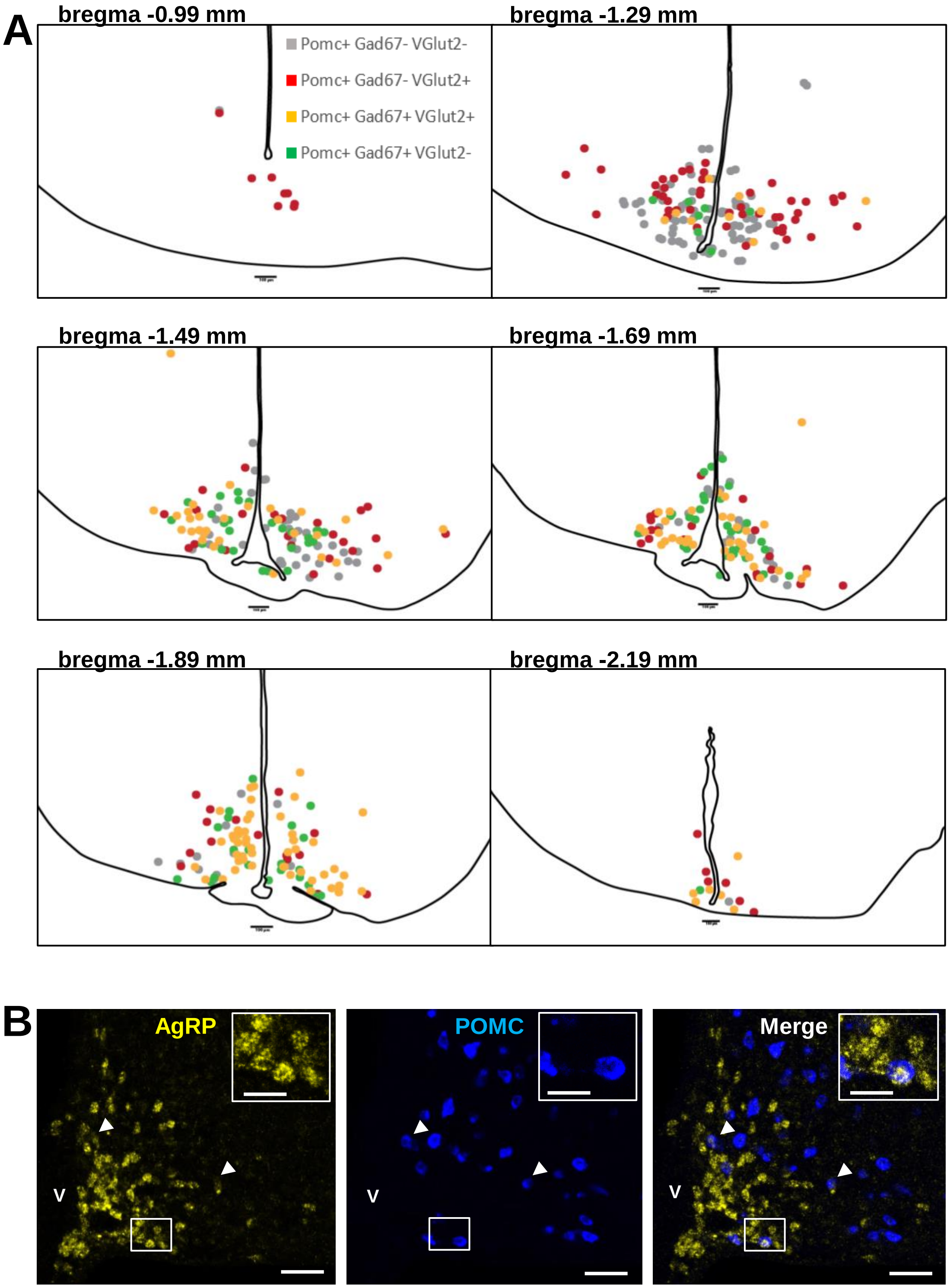

#### Figure S2, related to Figure 1

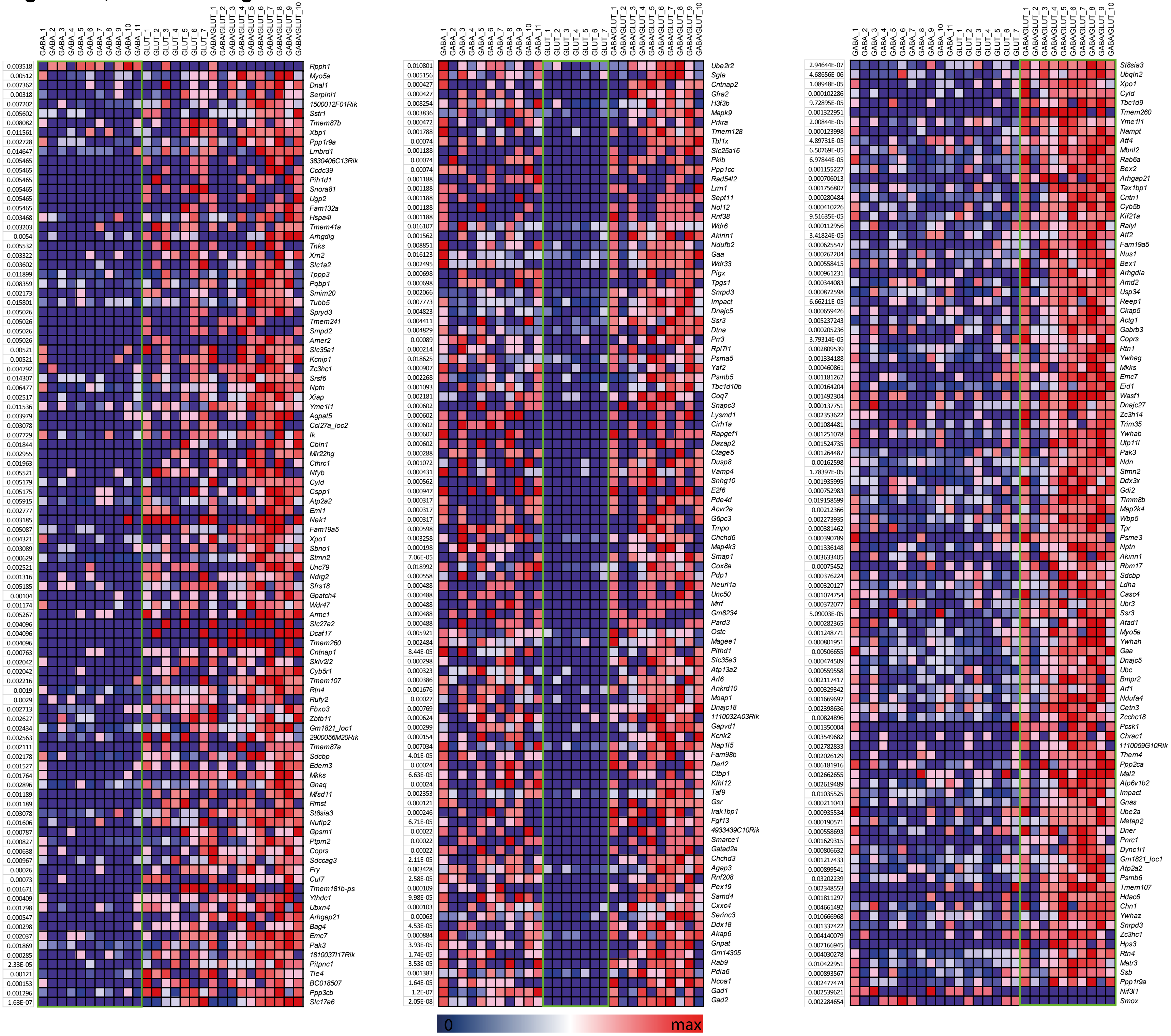

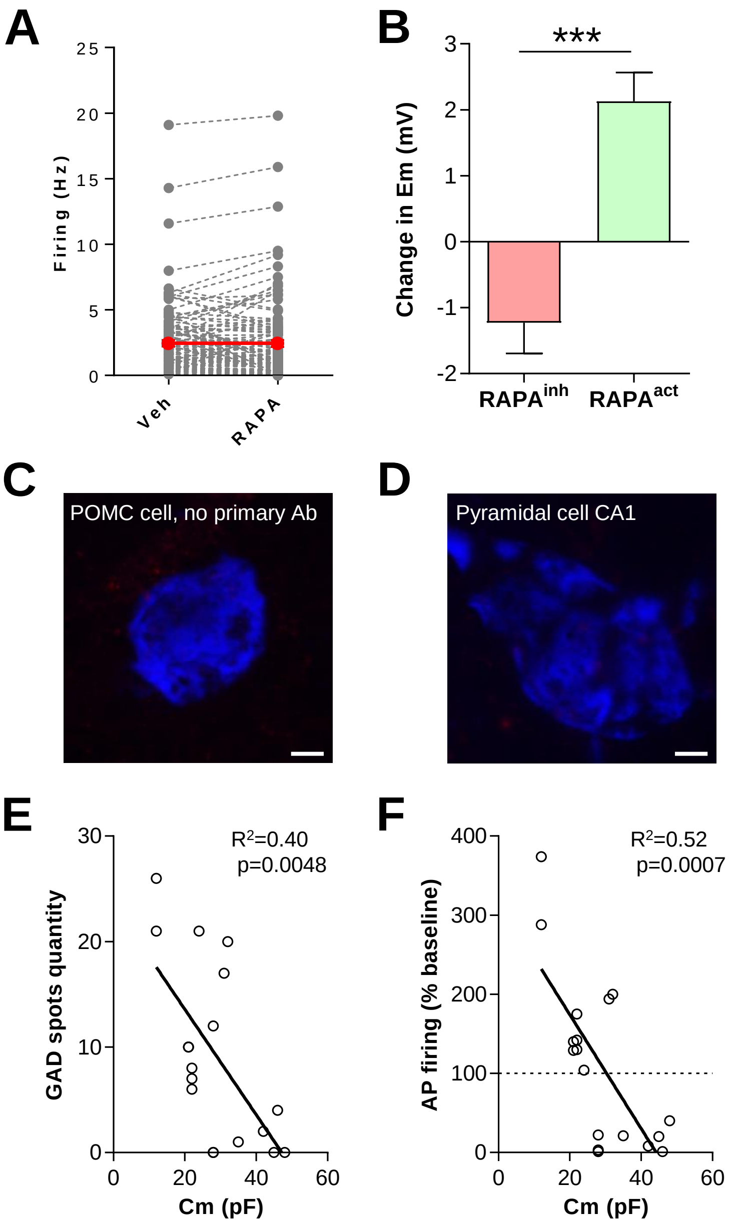

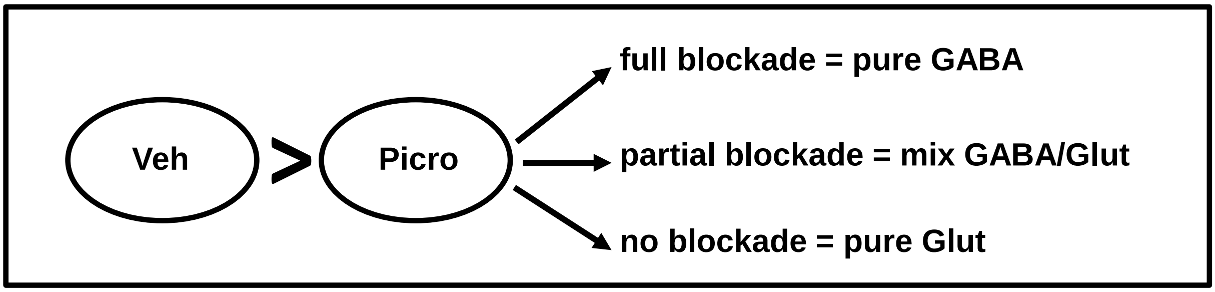

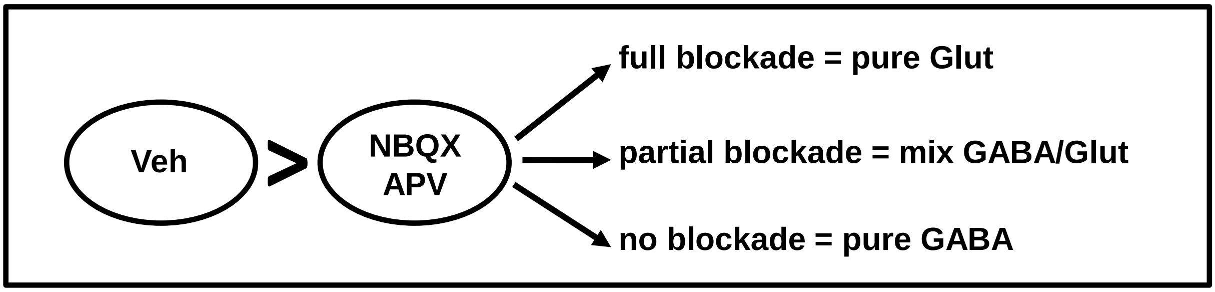

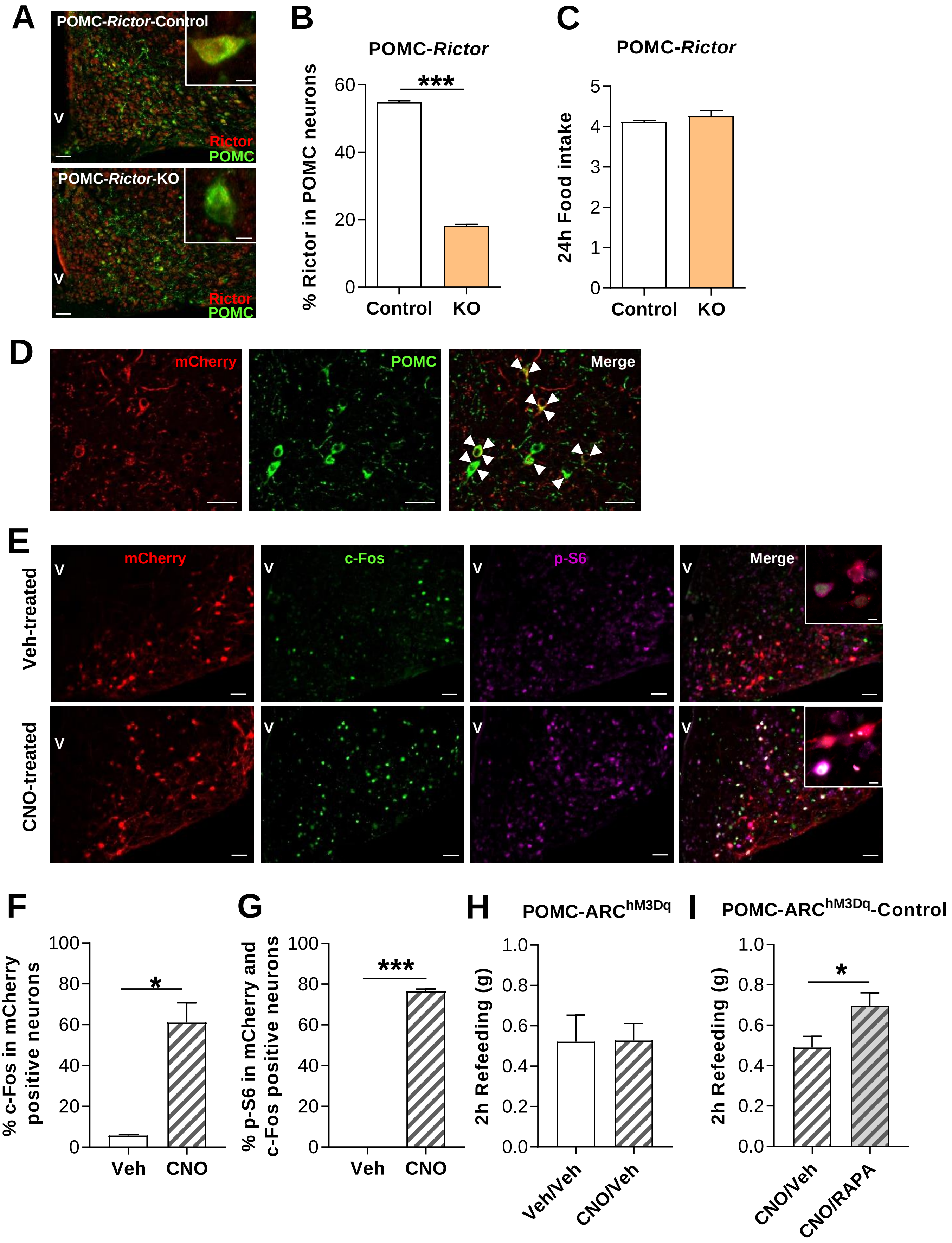

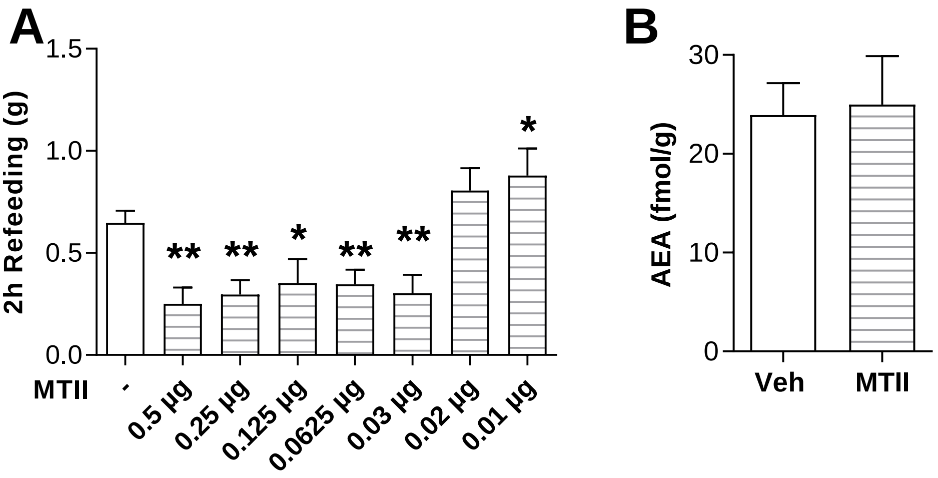

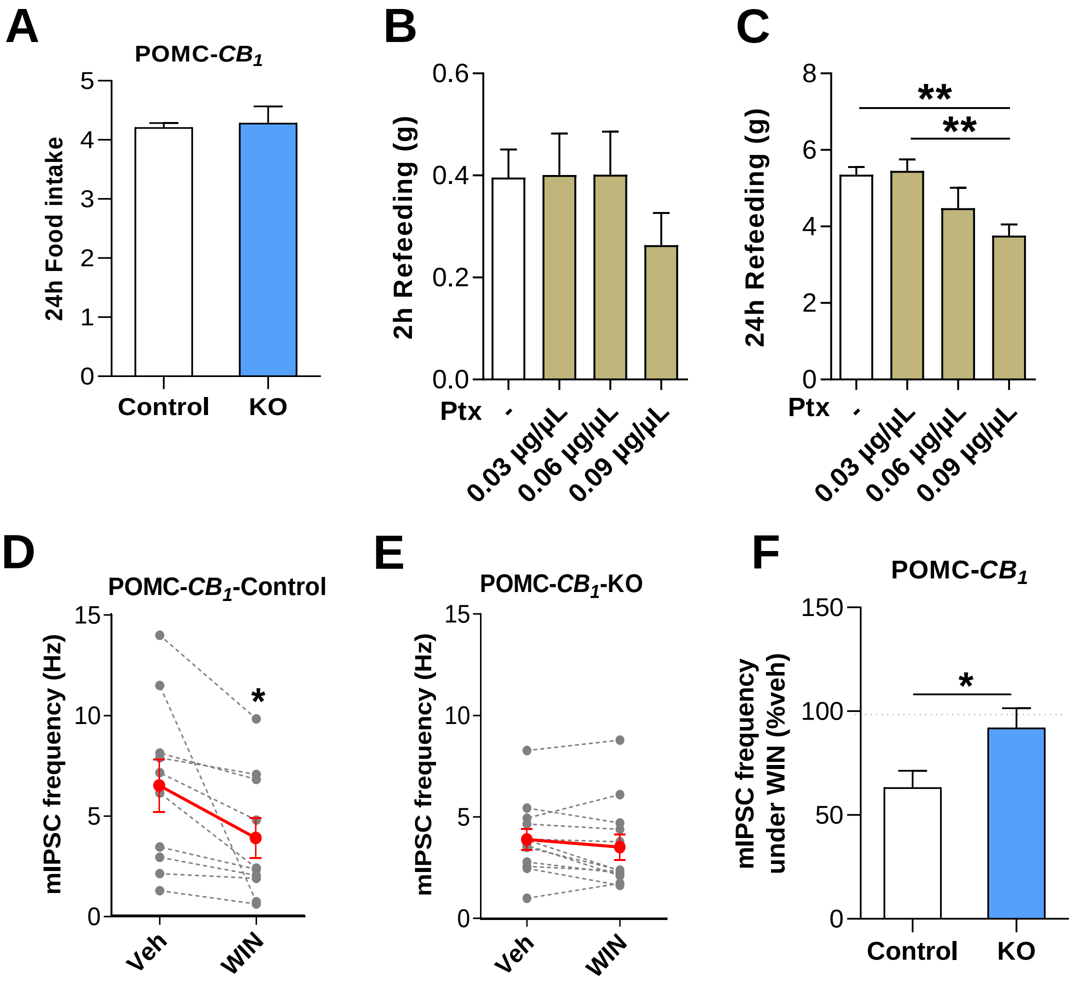

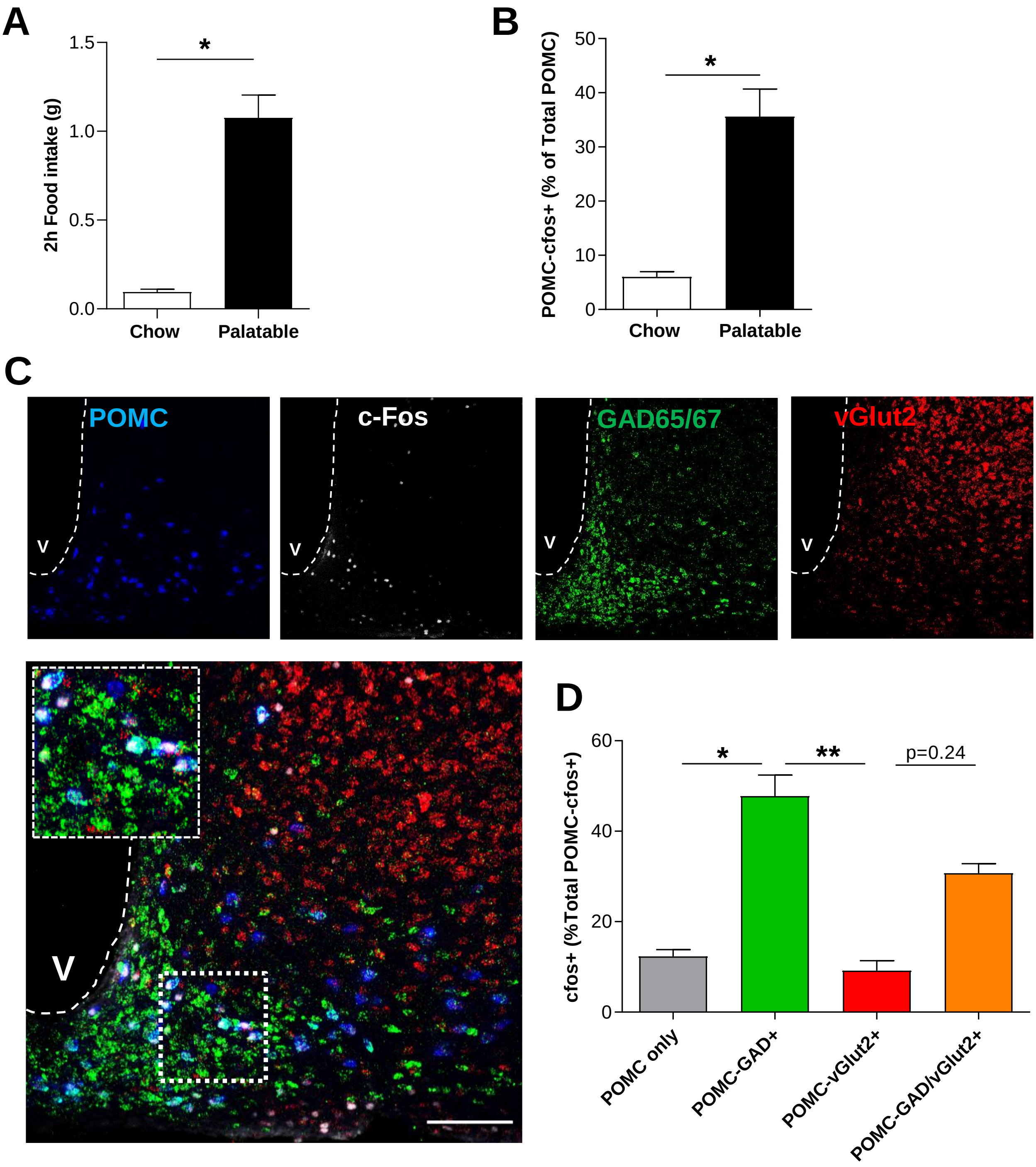

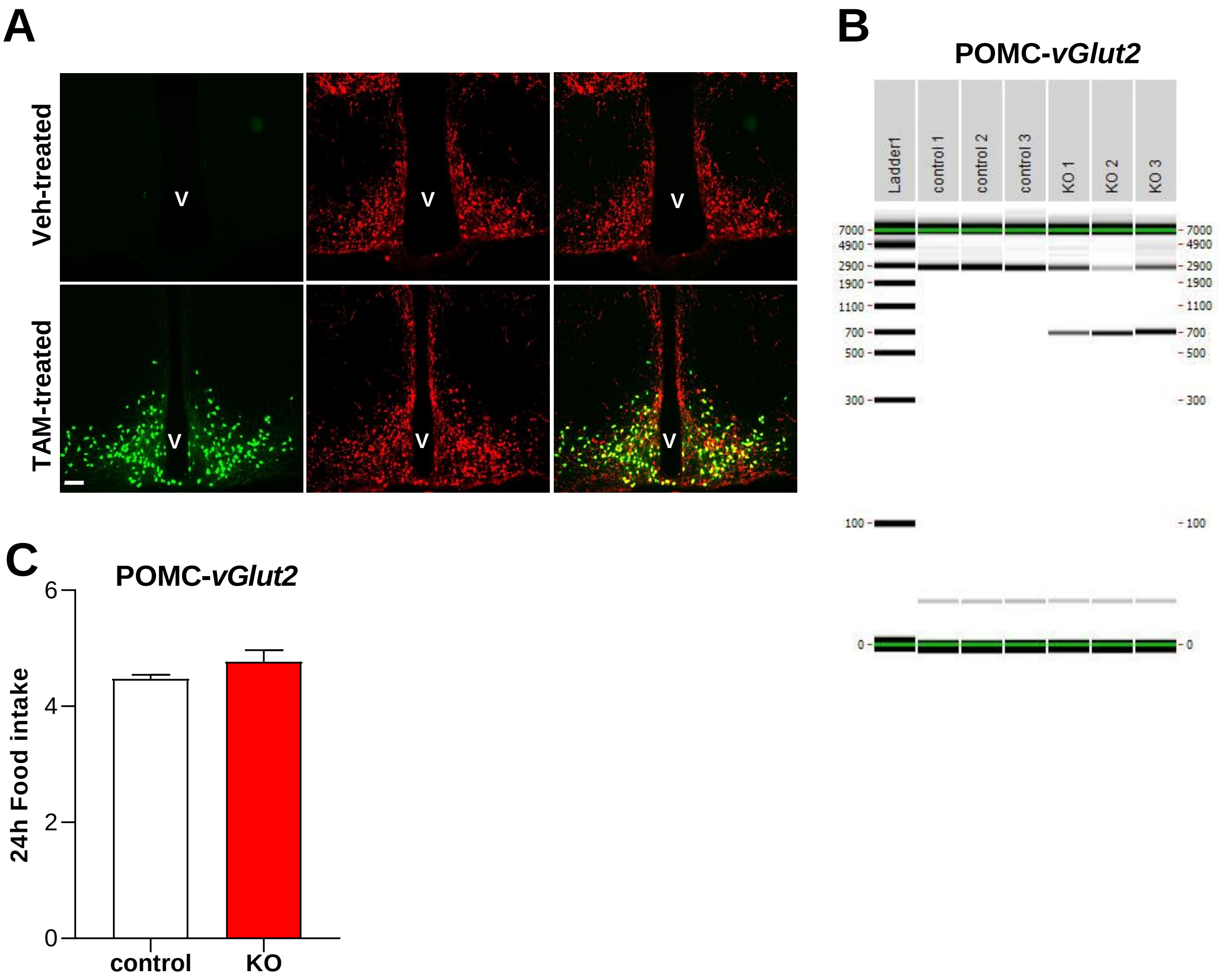
